# Comprehensive profiling of translation initiation in influenza virus infected cells

**DOI:** 10.1101/326967

**Authors:** Heather M. Machkovech, Jesse D. Bloom, Arvind R. Subramaniam

## Abstract

Translation can initiate at alternate, non-canonical start codons in response to stressful stimuli in mammalian cells. Recent studies suggest that viral infection and anti-viral responses alter sites of translation initiation, and in some cases, lead to production of novel immune epitopes. Here we systematically investigate the extent and impact of alternate translation initiation in cells infected with influenza virus. We perform evolutionary analyses that suggest selection against non-canonical initiation at CUG codons in influenza virus lineages that have adapted to mammalian hosts. We then use ribosome profiling with the initiation inhibitor lactidomycin to experimentally delineate translation initiation sites in a human lung epithelial cell line infected with influenza virus. We identify several candidate sites of alternate initiation in influenza mRNAs, all of which occur at AUG codons that are downstream of canonical initiation codons. One of these candidate downstream start sites truncates 14 amino acids from the N-terminus of the N1 neuraminidase protein, resulting in loss of its cytoplasmic tail and a portion of the transmembrane domain. This truncated neuraminidase protein is expressed on the cell surface during influenza virus infection, is enzymatically active, and is conserved in most N1 viral lineages. Host transcripts induced by the anti-viral response are enriched for translation initiation at non-canonical start sites and non-AUG start codons. Together, our results systematically map the landscape of translation initiation during influenza virus infection, and shed light on the evolutionary forces shaping this landscape.

## Author summary

When viruses such as influenza infect cells, both host and viral mRNAs are translated into proteins. Here we investigate the sites in these mRNAs that initiate protein translation during influenza infection. In particular, we explore whether some of this translation initiates at codons other than the canonical ones used to produce the primary protein product of each gene. Using computational analyses, we find that mammalian influenza viruses evolve to reduce the number of codons that can initiate such alternate translation initiation products. We next use the comprehensive experimental strategy of ribosome profiling to identify sites of translation initiation across all influenza and host mRNAs. We find a number of sites of alternate initiation on both influenza and host mRNAs. We study in detail one such alternate start site on an influenza mRNA, and show that it encodes a functional and previously uncharacterized variant of a viral protein. We also find evidence that the mRNAs that host cells express at higher levels during viral infection are enriched for translation initiation at non-canonical start codons. Overall, these results suggest that alternate translation initiation plays a role in shaping the repertoires of both viral and host proteins that are produced during influenza infection.

## Introduction

Cellular stress can impact protein translation initiation. For example, viral infection can activate protein kinase R, which globally reduces translation initiation for most transcripts, while translationally upregulating a subset of stress-response transcripts [1]. Cellular stress can also alter the start codon on mRNAs at which ribosomes initiate translation [2–4]. A common alternate start codon, CUG, may be of particular relevance during immune stress. Previous work using sensitive reporter assays found that CUG initiation can be induced during viral infection and activation of the immune response [5,6]. It has been hypothesized that upregulation of CUG initiation may be a deliberate host strategy to generate immune epitopes during cellular stress [5–7]. Indeed, there are numerous examples of immune epitopes that are initiated from alternate start sites, including CUG codons [8–13]. However, it is unknown if start site usage is altered globally during viral infection, or skewed towards certain start codons such as CUG.

In addition, viruses often rely on alternate translation initiation to increase the diversity of proteins encoded in their compact genomes. Influenza is an eight segmented negative stranded RNA virus that encodes ten canonical proteins, but additional influenza proteins can be generated through alternate initiation. For instance, the PB1-F2 protein is initiated at a downstream AUG in the +1 reading frame of the PB1 segment, and is thought to modulate the host immune response [14–16]. Additional peptides generated by alternate downstream AUG codons have been noted in the PB1, PA, and M influenza segments [14,15,17–21], but their abundance and functional relevance is unclear. There is some experimental support for CUG initiation in the non-coding region of M1 in the influenza genome [22], and very low levels of initiation on the negative sense RNA of the NS segment [23–25]. Even when such noncanonical peptides are generated at low levels and not functionally important for the virus, they can still be targeted by the host immune response [11]. However, despite the ad hoc identification of the aforementioned alternate protein products, there have been no systematic studies to map sites of translation initiation in all influenza transcripts.

Ribosome profiling, the high-throughput sequencing of ribosome-protected mRNA fragments, enables genome-wide interrogation of translation [26]. This method has been applied to influenza virus to reveal host shut-off [27] and translation of non-coding RNAs [28]. It has also been used to identify unannotated proteins in other viruses such as CMV [29]. Notably, ribosome profiling, in combination with inhibitors of translation initiation, can be used to identify candidate sites of translation initiation [30,31].

Here we integrate computational and experimental approaches to systematically examine translation initiation in influenza-infected cells. We first perform computational analyses that suggest there is selection against putative CUG alternate translation initiation sites in mammalian influenza virus lineages. We then use ribosome profiling to experimentally map sites of translation initiation in viral and cellular mRNAs during influenza virus infection. Surprisingly, we find little signal for CUG initiation in viral genes, but we do identify a number of candidate AUG start sites that are downstream of the canonical start codon. We show that one of these candidate start sites leads to production of a truncated version of the viral neuraminidase protein that is expressed on the cell surface and is enzymatically active. On the host side, we identify an excess of non-AUG start codons on transcripts that are upregulated during the anti-viral response. Together, our results highlight how alternate translation initiation can alter the protein-coding capacity of both viruses and the cells that they infect.

## Results

### Evolutionary analyses suggest selection against CUG start codons during influenza adaptation to mammalian hosts

We first examined whether we could identify any bioinformatic signature of evolutionary selection against alternate translation initiation. Such selection would occur if alternate translation initiation products are deleterious to the virus. One reason that such products might be deleterious is because they can serve as CD8 T-cell epitopes [8–13]. Human influenza virus evolves under pressure to escape CD8 T-cell immunity [32–34], so we hypothesized that the virus might minimize the number of alternate translation initiation sites as it adapts to humans.

We focused our bioinformatic analysis on alternate translation initiation at CUG codons for multiple reasons. First, several reporter studies have shown that CUG can serve as an alternate translation initiation codon [5,22,35–37]. Second, genome-wide ribosome profiling studies suggest that CUG is the most common non-AUG start codon [30,31,38]. Third, CUG-initiated peptides can serve as CD8 T-cell epitopes [5,37,39–41]. Finally, recent evidence suggests that immune responses and viral infection cause specific upregulation of CUG initiation [5,6].

One way to identify signatures of selection during influenza virus evolution is to compare changes in the genomes of viral lineages that have adapted to different host species [42,43]. Such host-specific adaptation occurs frequently when the virus jumps from its natural reservoir—wild waterfowl—to other host species including humans and swine [44]. We speculated that these host jumps could lead to selection against CD8 T-cell epitopes. Because humans are long lived and accumulate more adaptive immune memory, we further expected that immune selection on influenza will be generally stronger in humans than in birds and swine [45–50]

For our analysis, we examined five sets of influenza virus sequences. The first set includes human viruses descended from the 1918 pandemic strain, which is believed to have originated from an avian virus that jumped to humans [51–53]. Five of the eight segments in contemporary human H3N2 influenza virus (PB2, PA, NP, M, and NS) are descended in an unbroken lineage from the 1918 virus, and so have been adapting to humans for a century. We analyzed the protein-coding genes on these five segments. The second set includes the same genes from the classical lineage of swine influenza, which is thought to have also arisen from an avian viral strain around 1918 [51–53]. Our third and fourth sets comprise these genes from viruses that were isolated from ducks and chickens, respectively. Since birds represent the virus’s natural reservoir [54], we expect these viruses to be relatively stably adapted to their hosts. Our fifth set is sequences from human H5N1 infections, which are zoonotic infections by avian viruses that have had minimal time to adapt to humans [55].

To identify bioinformatic signatures of selection against alternate translation initiation, we began by examining whether there was depletion of CUG codons in reading frame 0 (in-frame with the canonical AUG) in human influenza. We found that the number of CUG codons in reading frame 0 of the human influenza virus lineage decreased over time (Fig. 1A, upper panel), with the recent human H3N2 viruses having about 40% fewer in-frame CUG codons than the 1918 virus. We also saw a similar trend in the swine lineage, suggesting that the loss of CUG codons results from a mammalian-specific rather than a human-specific evolutionary pressure. There is no notable decrease in the number of CUG codons in the avian viral sequences, which consistently have an in-frame CUG content similar to that of the 1918 virus. The in-frame CUG content of sequences from human H5N1 viruses (which mostly result from one-off infections with avian viruses) is similar to that of the avian and 1918 sequences. Notably, we did not see a comparable decrease in frequency of CUH (H = A/C/U) leucine codons in any of the five lineages (Fig. 1A, lower panel), indicating that the depletion of in-frame CUG codons is not attributable to a general selection against CUN leucine codons. This depletion is also not due to selection against the amino acid leucine or biochemically similar amino acids (Fig. S1). The depletion against CUG codons is still observed if we correct for each sequence’s nucleotide composition (Fig. S2).

**Fig 1.**
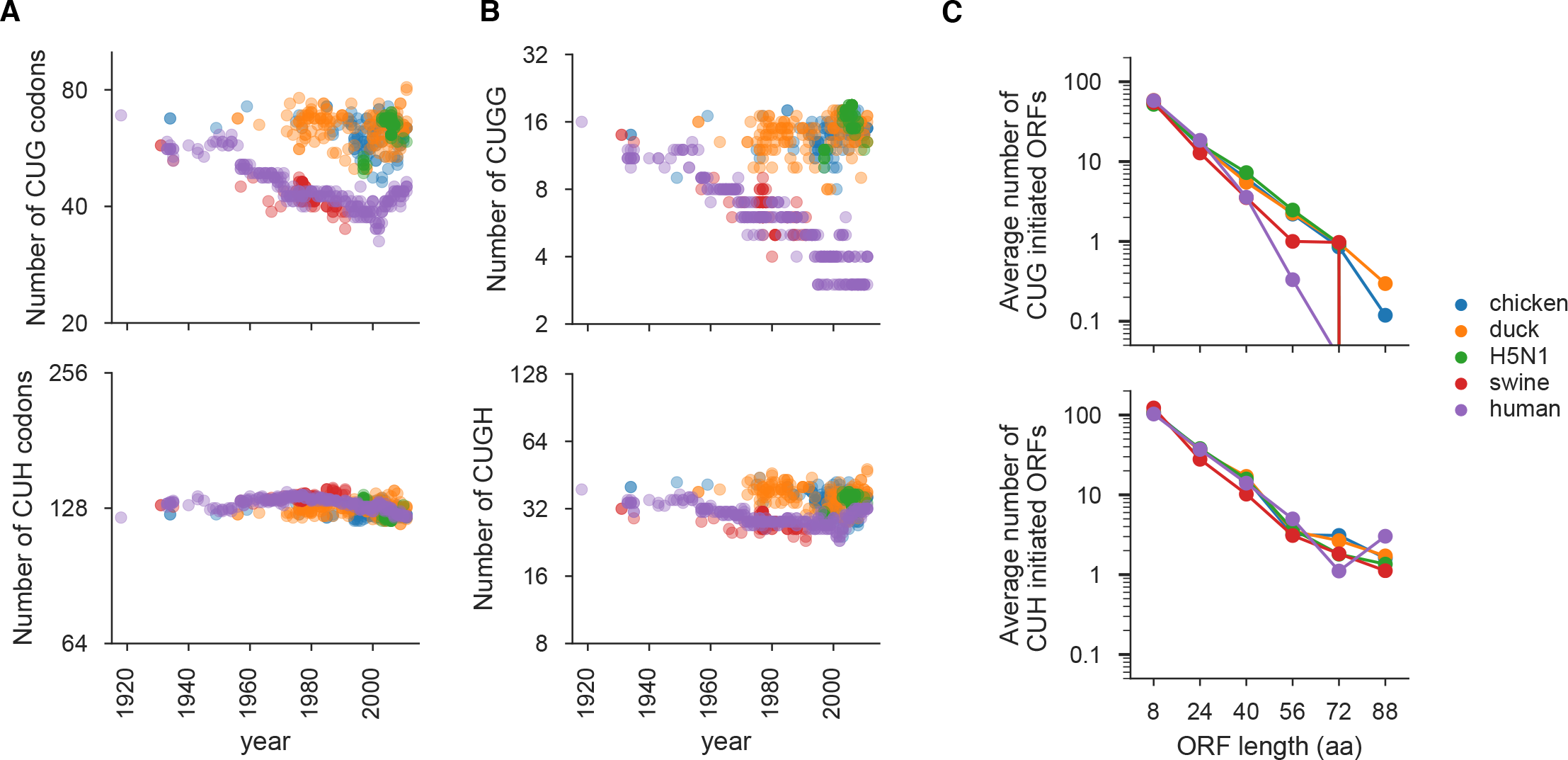
Evolutionary selection against CUG codons in mammalian influenza. **(A)** Number of CUG (upper panel) and CUH (where H is A, U or C, lower panel) codons in reading frame 0 of viral genes from human, avian, and swine influenza sequences. X-axis indicates the year that the viral sequence was isolated. **(B)** Number of in-frame CUGG (upper panel) and CUGH (lower panel) motifs in reading frame 0 of the viral genes. **(C)** Average length in codons of CUG- and CUH-initiated ORFs in reading frames 1 and 2 of the viral genes for sequences isolated between 1970 and 2011. All plots show data only for the genes on the PB2, PA, NP, NS and M segments, since these are the segments that have not re-assorted in human lineages derived from the 1918 virus.

Since the sequence context of a start codon plays an important role in determining how efficiently it can mediate translation initiation, we examined whether there is especially strong selection against CUG codons in sequence contexts that favor translation initiation. For AUG start codons, a purine at position −3 and a G at +4 can increase efficiency of initiation [56]. A recent reporter study has shown that CUG-mediated translation initiation is also promoted by a G at +4 [36]. Consistent with this study, we found that CUG initiation sites identified in mammalian transcripts using ribosome profiling are enriched for G at the +4 position (Fig. S3). Given the importance of the G at the +4 position, we examined whether there were differences in the depletion of in-frame CUGG and CUGH motifs in influenza virus. Indeed, we found that there is a more pronounced decrease in CUGG motifs than CUGH motifs in both human and swine viral lineages (Fig. 1B).

We next examined whether there was also selection against CUG codons that occur out-of-frame with the canonical AUG codon, as initiation in reading frames 1 and 2 would also generate novel peptides. We did not find evidence for selection against CUG codons in reading frames 1 and 2 (Fig. S2). However, in reading frames 1 and 2, eliminating the start codon is not the only way to eliminate long peptides initiated by alternate translation—these peptides can also be eliminated by introducing an early stop codon that shortens the peptide initiated by the alternate start codon. We therefore examined the length of putative CUG-initiated ORFs in reading frames 1 and 2. The human and swine viral lineages are depleted of long putative CUG-initiated ORFs (Fig. 1C, upper panel). Again, this selection is specific for CUG, since there is no host-specific depletion in long putative CUH-initiated ORFs (Fig. 1C, lower panel).

Since the viral sequences used in our analyses are phylogenetically related, we cannot easily test for the statistical significance of these evolutionary signals [57]. Nevertheless, the above analyses reveal bioinformatic signals consistent with selection against CUG-initiated alternate translation products in influenza virus lineages that have evolved in humans and swine.

### Systematic experimental profiling of translation initiation during influenza virus infection

Having established that human-adapted lineages of influenza virus exhibit signatures consistent with selection against alternate initiation at CUG codons, we sought to experimentally delineate the codons in influenza transcripts that could serve as potential sites of translation initiation during infection of a human lung epithelial cell line (A549). To this end, we employed ribosome profiling [26], a deep sequencing method for genome-wide quantification of ribosome-protected mRNA fragments. We performed two different variations of the ribosome profiling assay (Fig. 2A). We used the standard ribosome profiling method (Ribo-seq) to capture both elongating and initiating ribosomes. We did not add the elongation inhibitor cycloheximide to cells, since there can be artifacts associated with cycloheximide treatment [58]. We also performed a variation of ribosome profiling that enriches for initiating ribosomes [31] by pre-treating cells for 30 minutes with lactidomycin, a drug that preferentially binds empty E-sites of initiating ribosomes [59] thus allowing elongating ribosomes to run off mRNAs. Finally, we performed RNA-seq to profile the genome-wide transcriptional changes that occur during influenza infection.

**Fig 2.**
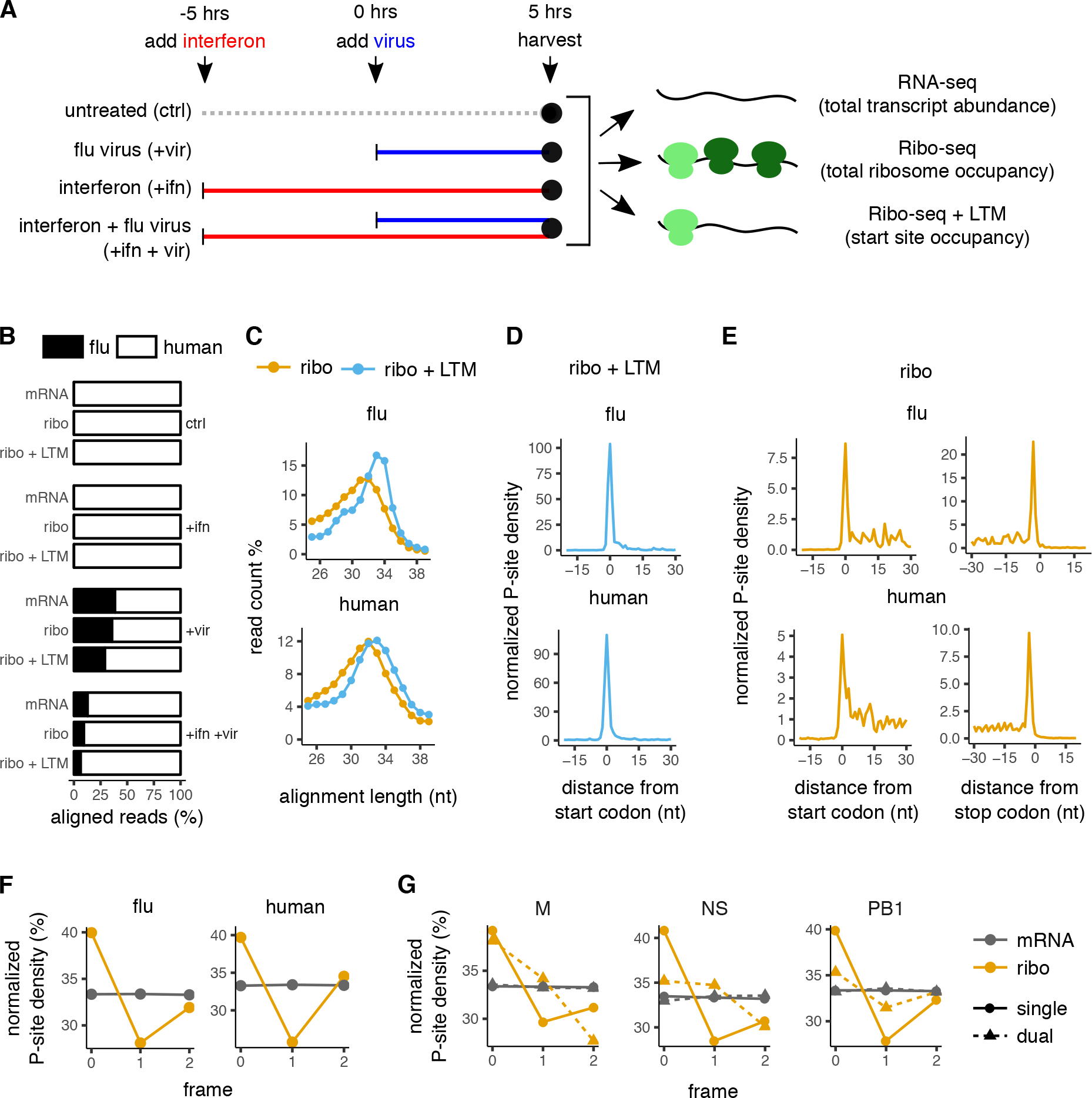
Translation initiation profiling of influenza infected cells. **(A)** Schematic of assays performed on A549 cells. Infections contained a mixture of viruses with their NP gene recoded to have low and high numbers of CUG codons (see Fig. S4). **(B)** Proportion of reads that align to viral and human transcripts for RNA-seq (mRNA), ribosome profiling (Ribo-seq), or Ribo-seq with LTM treatment. **(C)** Length distribution of reads that aligned to viral and human transcripts for Ribo-seq and Ribo-seq + LTM samples. **(D)** Metagene alignment of average P-site density around annotated start codons in viral and human transcripts for Ribo-seq + LTM sample. **(E)** Metagene alignment of average P-site density around annotated start and stop codons in viral and human transcripts for Ribo-seq samples. **(F)** Normalized P-site density in each of the reading frames of viral and human transcripts for RNA-seq (gray) and Ribo-seq (orange) samples. Both single- and known dual-coded regions in influenza transcripts were considered. **(G)** Normalized P-site density in each of the reading frames of single- and dual-coded regions of M, NS, and PB1 segments of the influenza genome for RNA-seq and Ribo-seq samples. Panels C–G show the +vir sample; see Fig. S5 for comparable plots for other samples. Only reads mapping to + sense of influenza transcripts are considered for panels B–G.

Anti-viral pathways have been shown to induce alternate translation initiation at CUG codons in reporter assays [6]. Therefore we also examined whether antiviral responses induced by interferon-*β* pretreatment alters the landscape of translation initiation in influenza virus. Altogether, we performed Ribo-seq and RNA-seq assays on four different samples of A549 cells (Fig. 2A): untreated cells (ctrl), interferon-*β* treated cells (+ifn), influenza virus infected cells (+vir), and cells that were pre-treated with interferon-*β* and subsequently infected with influenza virus (+ifn +vir). We utilized a reassortment virus with the polymerase components from A/PuertoRico/1934 and the remaining viral segments from A/WSN/1933. We used a high multiplicity of infection (MOI) such that most cells were productively infected by virus. We harvested cells 5 hours after viral infection, at which time the virus is expected to be amply transcribing and translating its mRNAs, but will not yet have undergone secondary rounds of infection [60,61].

To detect potentially low levels of alternate translation initiation at CUG codons in influenza transcripts, we used a mix of two viruses that were synonymously recoded in their NP gene to either deplete the number of in-frame CUG codons (low CUG NP), or
5/53 to increase the number of such codons (high CUG NP) (Fig. S4). Our rationale for recoding CUG codons in this gene is that NP is a major source of CD8 T-cell epitopes [62,63], and previous work suggested that CUG codons may initiate peptides that contribute to the immune epitope pool [5,6]. Using this virus mix allowed us to have an internally controlled experiment: if a CUG codon in the high CUG NP variant can initiate translation, we should be able to preferentially detect it against the low CUG NP variant background even in the presence of end-specific sequence biases in the ribosome profiling method [64].

Deep sequencing of ribosome protected fragments yielded 15–40 million reads, of which 2–3 million reads could be mapped to human or influenza transcripts after standard preprocessing steps (Fig. S5A). As observed previously [27], 15–25% of influenza-aligned reads mapped to the negative sense influenza genomic RNAs (Fig. S5A). In contrast to the reads mapping to the positive sense strand of influenza (Fig. 2C), the negative sense reads did not show a characteristic peak around the size of a ribosome footprint (Fig. S5D), and likely arise from co-purification of influenza genomic RNAs that are protected from nuclease digestion due to their association with the viral NP protein [65,66]. Therefore, we only considered reads that mapped to the positive sense strand of influenza transcripts for the remaining analyses. Reads mapping to influenza transcripts accounted for between 29–39% of the mapped reads in the virus-infected samples (Fig. 2B), consistent with our use of high MOI. As expected, interferon-*β* pretreatment reduced productive infection, and reads mapping to influenza transcripts only accounted for 6-13% of aligned reads in the +ifn +vir sample (Fig. 2B).

We performed additional checks to ensure adequate quality of our Ribo-seq data. Ribo-seq reads mapping to either human and influenza transcripts had read lengths between 29–34 nucleotides (Fig. 2C, Fig. S5B). This length distribution was on average a few nucleotides wider and longer than observed in other ribosome profiling studies [30,67,68], likely because of less stringent RNase I digestion. We assigned the ribosome P-site to either the 14th or 15th nucleotide from the 5′ end of footprint reads depending on their length (see Methods). Consistent with previous observations [31], mean P-site density in the Ribo-seq + LTM samples showed >100-fold enrichment at the annotated start codons of both human and influenza transcripts (Fig. 2D, Fig. S5C). P-site density in the Ribo-seq samples exhibited a peak at start and stop codons and was distributed within the coding region of human and influenza transcripts (Fig. 2E, Fig. S5C).

The mean P-site density in Ribo-seq samples displayed the expected 3-nucleotide periodicity along human transcripts (Fig. S5E), and was enriched in frame 0 of both human and influenza coding sequences (Fig. 2F, S5F). Notably, the known 191 nucleotide dual-coding region in the NS segment of the influenza genome [69] had a distinct P-site density distribution across the three reading frames compared to the single-coding region of the same segment (Fig. 2G, S5G). This dual-coding signature was present but less pronounced in the known dual-coding regions of M (45 nucleotides) and PB1 (261 nucleotides) segments (Fig. 2G, S5G), likely because M2 and PB1-F2 are expressed at lower levels during infection [70–72].

Finally, we used our sequencing data to evaluate the quality of the virus stocks used in our experiment. Virus stocks contain virions that range in biological activity, including virions defective in replication (defective particles) [73–77]. For influenza virus, defective particles often contain large internal deletions in the polymerase segments [73–78]. The presence of defective particles with internal deletions would diminish our ability to detect alternate initiation sites within the deleted regions. However, the burden of defective viral particles can be reduced by growing virus for a short amount of time and at a low MOI, as defective viral particles increase in frequency when viruses are grown at a high MOI where complementation of deleterious genotypes can occur [79]. The even read coverage across the polymerase segments in our data (Fig. S6B) indicates that we had a low burden of defective particles. This low burden was particularly evident when we compared our data to a previous ribosome profiling study of influenza virus infected cells (Fig. S6A) [27].

### Translation initiation sites in the viral genome

We next used the Ribo-seq and Ribo-seq + LTM measurements to annotate candidate translation initiation sites (TIS) in the influenza genome. Our general strategy to identify candidate TIS was to find peaks in Ribo-seq + LTM coverage within each influenza transcript that was significantly higher than both the Ribo-seq coverage at the corresponding location and the background Ribo-seq + LTM coverage distribution for that transcript (Fig. 3A). Specifically, we used a zero-truncated negative binomial distribution (ZTNB) to statistically model the background distribution of Ribo-seq and Ribo-seq + LTM counts in each transcript [80,81]. Candidate start sites were identified based on the following criteria: The ZTNB-based P-value for the Ribo-seq + LTM count at that location must be <0.01 and 1000-fold higher than the P-value of the Ribo-seq counts at the same location (Fig. 3A, left panel), or must have an absolute value less than 10^−7^. In addition, the Ribo-seq + LTM counts must be a local maximum within a 30 nucleotide window (Fig. 3A, right panel), and the Ribo-seq counts must be non-zero. To account for the 1–2 nucleotide positional uncertainty of the P-site density in our Ribo-seq and Ribo-seq + LTM measurements, we pooled the read counts in 3 nucleotide windows before applying the above criteria.

**Fig 3.**
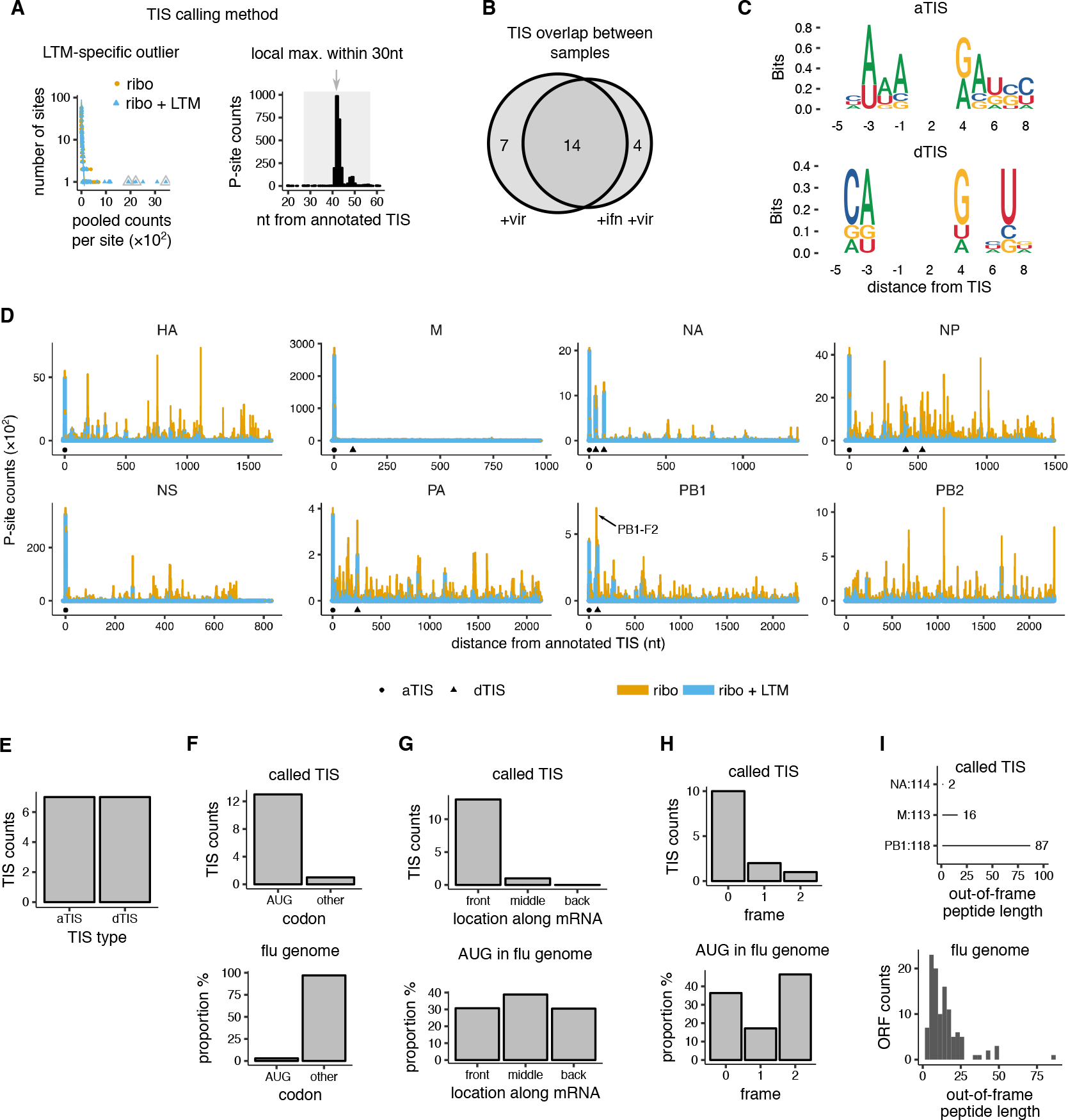
Candidate translation initiation sites on influenza transcripts. **(A)** Method for identifying candidate translation initiation sites (TIS) in influenza. Left panel: LTM-specific outliers were identified over the background of Ribo-seq and Ribo-seq + LTM pooled counts at each site of each influenza transcript. Right-panel: Among LTM-specific outliers, only the highest LTM peak in P-site counts within each 30 nucleotide window was called as start site. The background Ribo-seq and Ribo-seq + LTM counts were fit to separate zero-truncated negative binomial distributions (lines in left panel). The final called TIS are indicated by grey triangles and arrows. The left panel shows the NA segment of our +vir sample and the right panel a window around the NA43 TIS. See Fig. S7 for other influenza segments. **(B)** Overlap in candidate TIS between the +vir and +ifn +vir samples. **(C)** Consensus motif of annotated translation initiation sites (aTIS, upper panel) and downstream translation initiation sites (dTIS, lower panel). **(D)** P-site counts from Ribo-seq and Ribo-seq + LTM assays are shown for all 8 influenza genome segments for our +vir sample. The counts from the two assays are shown as stacked bar graphs for ease of comparison. The candidate annotated TIS (circle) and downstream TIS (triangle) are indicated below the coverage plots. The known alternate initiation site PB1-F2 is highlighted with arrow. **(E)** Number of called annotated and downstream TIS. **(F)** Distribution of codon identity for candidate influenza TIS (top) and for all codons in influenza genome (bottom). Other indicates all non-AUG codons. **(G)** Distribution of AUG codons along transcripts for candidate influenza TIS (top) and for all AUG codons (bottom). The codon locations were binned into three fragments of equal length for each influenza CDS. **(H)** Distribution among reading frames for candidate downstream TIS (top) and for all AUG codons in the influenza genome (bottom). **(I)** Length of peptides initiated by called out-of-frame TIS (top) and all out-of-frame AUG codons in the influenza genome (bottom). Only the candidate TIS shared between the +vir and +ifn +vir samples were used for panels C–I.

We applied our candidate TIS identification strategy to each influenza transcript (Fig. S7) separately for the +vir and +ifn +vir samples. We assigned the candidate TIS peaks to any near-cognate AUG codon (at most one mismatch from AUG) if that codon was within 1 nucleotide of either side of the TIS peak. We identified a total of 25 candidate TIS across both samples (Fig. S8). Fourteen of the 25 identified TIS overlapped between the two samples (Fig. 3B), and we used this overlapping subset as a high-confidence set of candidate TIS for downstream analyses. We did not detect a higher number of candidate TIS in our +ifn +vir sample compared to the +vir sample (4 vs. 7, Fig. 3B), suggesting that anti-viral response as mediated by interferon-*β* induction does not result in detectably higher number of alternate translation initiation sites in influenza transcripts [6]. An important caveat to this observation is that the Ribo-seq + LTM assay is likely to miss TIS that have a low frequency of initiation, but could still be detectable by sensitive immunological assays [5,25].

The high-confidence set of 14 candidate TIS had multiple features consistent with being bona fide TIS. First, this set included 7 of the 8 annotated TIS (aTIS) for the 8 segments in the influenza genome (Fig. 3E). Only the annotated TIS of the PB2 segment was not identified, and this is due to the dense Ribo-seq + LTM coverage and lack of clear peaks in this segment (Fig. 3D, bottom right panel). Second, our set also correctly identified the start codon for two previously annotated protein products generated by initiation in an alternate reading frame of the PB1 and M genes. Initiation in the +1 reading frame at nucleotide 118 in the PB1 gene generates PB1-F2 (Fig. 3D, arrow). Initiation in the +1 reading frame at nucleotide 113 in the M segment is also a known to occur [21]. Third, 13 of the 14 candidate TIS had an AUG codon within 1 nucleotide even though less then 3% of trinucleotides in the influenza genome are AUG (Fig. 3F, top vs. bottom). Fourth, both the annotated and the novel candidate TIS in our set had an over-representation of A at −3 nucleotide and G at +4 nucleotide positions (Fig. 3C), which are known to be optimal contexts for translation initiation in vertebrate cells [56]. Finally, the candidate TIS are enriched towards the 5′ end of the transcripts, with 13 of the 14 candidate TIS located in the initial third of the influenza transcripts (Fig. 3G). This last observation is consistent with the initiation of candidate TIS through scanning by the 43S pre-initiation complex from the 5′ end [82].

Six of the 7 alternate TIS in our candidate set are AUG codons that are downstream of the respective canonical aTIS (Fig. 3E). This downstream location of alternate TIS is expected given the short 5′ UTR (around 20 nucleotides) of influenza transcripts. While previous translation initiation profiling studies have identified a large number of putative non-AUG TIS [30,31], these are predominantly located in the much longer 5′ UTRs of mammalian transcripts, upstream of aTIS. Even for mammalian transcripts, a majority of TIS identified downstream of annotated start codons are at AUG codons [83].

The alternate dTIS in our high-confidence set are distributed across multiple influenza segments: M, NA, NP, PB1, and PA (Fig. 3D, Fig S9). Ten of the 13 high-confidence TIS initiating at AUG are located in the canonical reading frame 0 (Fig. 3H). This set includes 7 annotated AUG starts, as well as three in-frame downstream AUGs in NA, NP, and PA segments that would result in N-terminally truncated forms of the annotated proteins. The three out-of-frame candidate dTIS are the known PB1-F2 ORF [14], the known start codon at nucleotide 113 of the M gene [21], and a short ORF of length 2 in NA (Fig. 3I, top panel). Excluding the well-characterized PB1-F2 ORF, the two candidate out-of-frame candidate ORFs have lengths that are typical of out-of-frame AUG-initiated sequences in the influenza genome (Fig. 3I, lower panel).

Among the other alternate translation initiation sites in the influenza genome that have been described previously, we did not find evidence for the initiation sites previously noted in PA and PB1 (see Fig. S10) [14,15,17–20]. This could be because of low initiation frequency at these sites under our infection conditions. Re-analysis of the harringtonine-treated ribosome profiling data from Ref. [27] revealed P-site count peaks that coincided with all the 7 annotated TIS as well as 3 of the 7 downstream TIS identified in our experiments (Fig. S11). Among the 4 dTIS that could not be clearly associated with a harringtonine peak, 3 are >200 nucleotide from the 5′ end of the transcripts, and could have been potentially affected by the high fraction of defective viral particles in [27] (Fig. S6A). Interestingly, the dTIS in our dataset without a corresponding harringtonine peak is an in-frame AUG in the NA segment that is mutated to the near-cognate CUG codon in the PR8 influenza strain used in [27].

### Translation initiation at CUG codons in influenza NP

As mentioned previously, our infections were performed with a mix of two otherwise isogenic viruses that were recoded to have different numbers of CUG codons in the NP gene (Fig. S4). This experimental design allows us to sensitively look for CUG initiation that cannot be detected by the start-site calling method in the previous section, but might still be detectable via an internally controlled comparison of the two NP variants.

Specifically, the high CUG NP variant has 20 leucine codons that were synonymously mutated to a non-CUG codon in the low CUG NP variant (Fig. 4A). We examined if there was evidence of enhanced initiation at any of these codon sites in the high CUG NP variant. Between 35% and 42% of reads mapping to NP could be uniquely assigned to either the high or low CUG NP sequences based on polymorphisms due to the synonymous mutations. These reads mapped to the two variants at a nearly equal (1:1.3 ratio) overall proportion (Fig. 4A). The ratio of P-site density between the two variants at individual NP sites obtained from the uniquely-mapping reads varied over a 500-fold range (Fig. 4B). Among the sites with an excess of high CUG NP P-site density, the CUG codon 322 nucleotides from the annotated TIS displayed the largest such excess with 16-fold more reads for the high CUG NP variant than the low CUG NP variant (Fig. 4B, red point). This excess P-site density was present in both the Ribo-seq and the Ribo-seq + LTM data (Fig. 4C). The length distribution of the reads generating this excess was consistent with them being derived from ribosome-protected fragments (Fig. S12). The excess P-site density at CUG322 did not arise from 3′ ends of reads mapping to the nearby recoded CUG328 (Fig. S13). The +ifn +vir sample displayed a similar excess of density at site 322 for the high CUG NP variant (Fig. S14).

**Fig 4.**
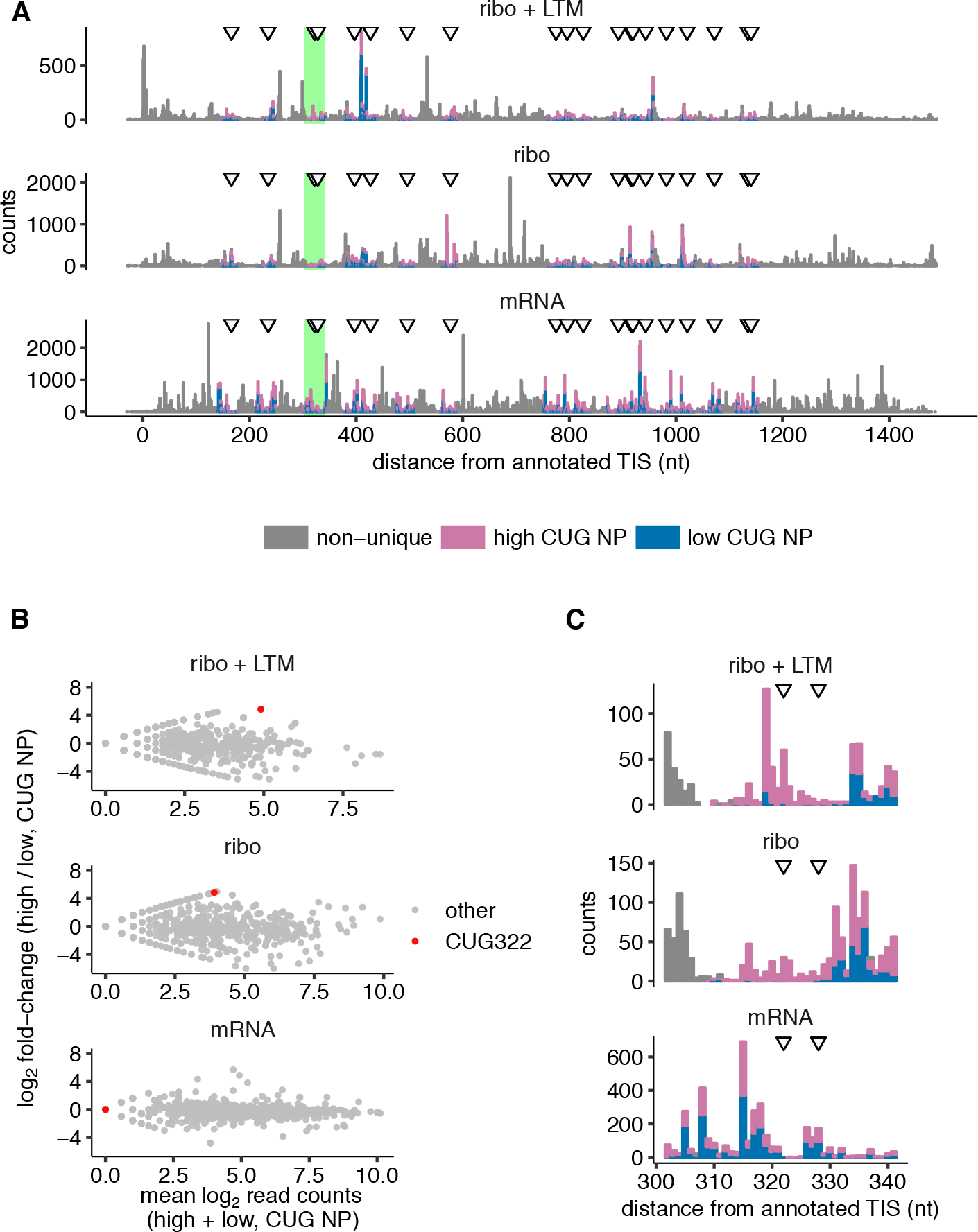
Translation initiation at CUG codons in influenza nucleoprotein. **(A)** Coverage of Ribo-seq + LTM, Ribo-seq, and RNA-seq reads that can be uniquely aligned to either the high CUG NP variant or the low CUG NP variant, along with reads that cannot be uniquely assigned. P-site counts are shown for Ribo-seq and Ribo-seq + LTM assays. 5’-end counts are shown for RNA-Seq. Data are plotted as a stacked bar graph. Locations of the 20 CUG codons that are present in high CUG NP and synonymously mutated in low CUG NP are indicated by arrows. **(B)** The ratio of high CUG NP to low CUG NP coverage from panel A is plotted against their sum along the horizontal axis. **(C)** The green-highlighted region in panel A around the CUG322 codon is shown at greater horizontal magnification. Data shown for +vir sample. See Fig. S14 for +ifn +vir sample.

To examine whether CUG322 could initiate translation in the high CUG NP variant, we used Western blots of heterologously expressed NP variants. However, we were unable to resolve any protein fragment of appropriate size that was present only in the high CUG NP variant (Fig. S15). It is possible that there is no initiation at this site, or that initiation occurs but at a very low level, consistent with the low overall Ribo-seq + LTM P-site density at CUG322 compared to the other locations on NP. In that case, more sensitive immunological assays [5,25] might be necessary to identify initiation at CUG322 and other CUG codons in the high CUG NP variant. Another important caveat is that LTM treatment may not effectively arrest CUG-initiating ribosomes. Indeed, evidence for CUG-based initiation being refractory to several translation inhibitors has been noted in previous reporter-based studies [5].

### Functional characterization of an in-frame alternate start site in NA

Among the three in-frame alternate TIS identified by our Ribo-seq experiments (Fig. 3D), the AUG at nucleotide 43 from the aTIS of the neuraminidase (NA) gene had the highest statistical significance in our start site calling method (Fig. S8). This candidate TIS (Fig. 5A) had several other features that prompted us to investigate it in further detail.

**Fig 5.**
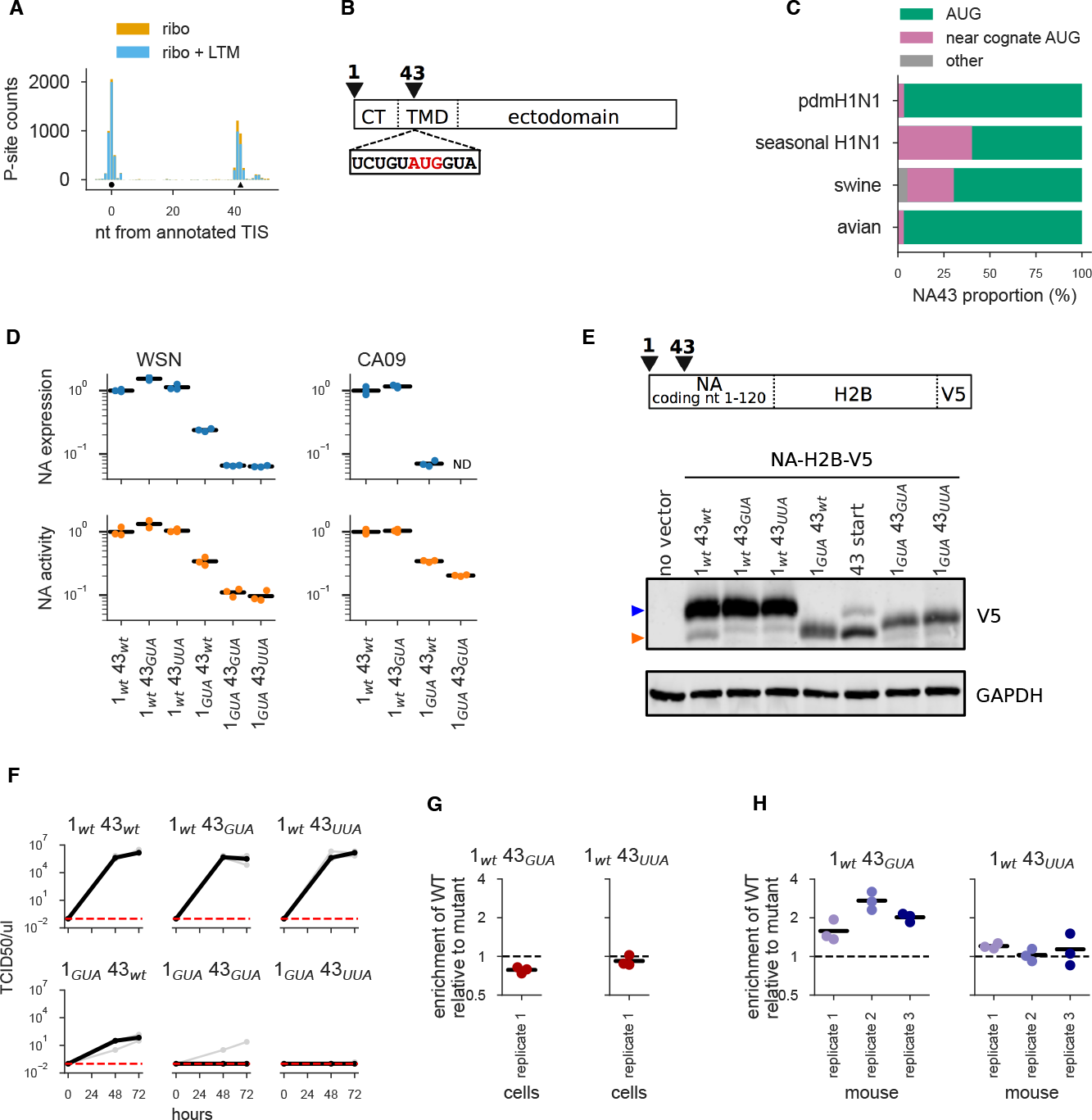
Characterization of the NA43 protein generated by alternate translation initiation in the NA gene. **(A)** P-site counts from Ribo-seq and Ribo-seq + LTM assays are shown as stacked bar graphs for the 5′ end of the mRNA for our +vir sample. The candidate annotated TIS (circle) at coding nucleotide position 1 and the downstream TIS (triangle) at coding nucleotide position 43 are indicated below the coverage plots. **(B)** Schematic of NA’s primary sequence. NA’s cytoplasmic tail (CT) spans amino acids 1-6, the transmembrane domain (TMD) spans amino acids 7-35, and the ectodomain spans amino acids 36 to 453. The inset shows the AUG43 codon (in red) and surrounding sequence. **(C)** Most NAs in major lineages of N1 influenza have either an AUG or a near-cognate start codon at nucleotide 43. **(D)** Quantification of NA cell-surface protein levels and enzymatic activity in 293T cells transfected with plasmids encoding NA variants with mutations at the canonical or downstream start site. Results are shown for NA from the WSN and CA09 viral strains. Measurements are normalized relative to the mean for wildtype, and black lines indicate the mean of three measurements for all variants except for WSN 1_wt_*43*_GUA_, which only had two measurements. ND indicates that the value was below the limit of detection. **(E)** Schematic of coding region of NA-H2B-V5 constructs used for Western blots. Blot of 293T cells transfected with NA-H2B-V5 constructs with mutations at the canonical or downstream start site. “43 start” is a size control construct that begins at site 43 of NA. Top panel: anti-V5 (blue arrow corresponds to full length NA and orange arrow corresponds to NA43); bottom panel: anti-GAPDH. **(F)** Viral titers in the supernatant at 48 and 72 hours post-transfection during reverse-genetics generation of WSN viruses carrying NA with the indicated mutations. Three independent replicates for each mutant are shown in gray, and the median is shown in black. Undetectable titers are plotted at the assay’s limit of detection of 0.1 TCID50 / *μ*l. **(G)** *In vitro* competition of wildtype virus versus virus carrying the indicated NA mutation to ablate the initiation codon at site 43. We mixed wildtype virus and mutant virus at a target ratio of 1:1 infectious particles, infected cells, and collected viral RNA at 10 hours and 72 hours post-infection to determine variant frequencies by deep sequencing. Shown is the enrichment of wildtype to mutant for the 72 hour timepoint relative to the 10 hour timepoint. The black bar indicates the mean of the three biological replicates. There was no consistent enrichment of the wildtype virus relative to the mutant lacking NA43. **(H)** Similar competition experiments performed *in vivo* in mice. We collected samples at 96 hours post-infection to determine viral frequencies. Shown is the enrichment of wildtype to mutant after 96 hours of viral growth in mice relative to the 10 hour cell culture timepoint. The black bar indicates the mean of the three biological replicates. We performed a paired *t*-test with the resulting *P*-values: 0.09 (*1_wt_43_GUA_*), 0.11 (*1_wt_43_UUA_*).

The AUG43 in NA has a favorable translation initiation context, with a G at +4 nucleotides (Fig. 5B). The function of NA is to mediate viral egress by cleaving the viral receptor sialic acid from the cell surface [84,85]. NA is a type II membrane protein [86,87], meaning that the cytoplasmic tail and the transmembrane domain are at the N-terminus and the ectodomain is at the C-terminus (Fig. 5B). The truncated NA protein resulting from translation initiation at AUG43 would lack the cytoplasmic tail and the first few amino acids of transmembrane domain. Interestingly, classical studies of type II membrane proteins characterized a series of artificial mutants of NA in an effort to determine the motifs required for membrane insertion of the protein [88,89]. One of these NA mutants was an N-terminal deletion in which the first 42 nucleotides were removed, effectively creating the NA43 protein that would be generated by translation initiation at AUG43. In the context of a protein expression plasmid, this NA N-terminal deletion was efficiently expressed and localized to the cell surface, indicating that the signal and anchor domains are reasonably intact [88,89]. On the basis of this prior work, we hypothesized that any NA43 protein generated by alternate translation initiation at AUG43 of the wildtype NA gene would also create a cell-surface NA protein lacking the cytoplasmic tail and a portion of the transmembrane domain.

We analyzed the sequences of NA genes across different influenza lineages to examine if the AUG43 codon is evolutionarily conserved. Most N1 avian influenza and human pdmH1N1 NAs have an AUG at nucleotide 43, as do the majority of N1 swine influenza NAs (Fig. 5C). Some human seasonal H1N1 NAs and N1 swine influenza NAs lack an AUG at nucleotide 43, but these strains usually have another near-cognate start codon at the site instead (Fig. 5C). Therefore, the candidate downstream start site at NA nucleotide 43 is present in most N1 influenza strains.

We next sought direct experimental evidence for the NA43 protein initiated by the downstream start site. Towards this, we generated a series of mutants of the NA from the WSN strain of influenza. In these constructs, we mutated one or both of the canonical and candidate downstream start site to codons that are not near-cognate AUG codons. *1_wt_43_wt_* contains the wildtype AUG codon at both the canonical and candidate downstream start sites, *1_GUA_43_wt_* has the canonical start codon mutated to GUA, *1_wt_43_GUA_* and *1_wt_43_UUA_* have the candidate downstream start site mutated to GUA or UUA, and *1_GUA_43_GUA_* and *1_GUA_43_UUA_* have both codons mutated to the indicated identities. If both the canonical and downstream start sites initiate translation, then these constructs should encode both, just one, or neither of the full-length NA and NA43 proteins.

We first used these constructs to test whether NA43 is produced and localized on the cell surface. We did this by transfecting 293T cells with protein expression plasmids encoding the various NA mutants with a C-terminal V5 tag, which does not disrupt NA folding or function and can be detected by flow cytometry [90]. NA protein was detected on the surface of cells transfected with plasmids encoding mutants that lacked an AUG at the canonical start codon (Fig. 5D, top left). Specifically, transfection of cells with the *1_GUA_43_wt_* mutant led to ~24% of the cell-surface NA found in cells transfected with wildtype NA. Production of this cell-surface NA in mutants lacking the canonical start codon is mostly dependent on the candidate downstream start site at nucleotide 43, since mutants that lack AUG at this position (*1_GUA_43_GUA_* and *1_GUA_43_UUA_*) had much lower levels of cell-surface NA (Fig. 5D, top left). Interestingly, when we mutated the candidate downstream start site but not the canonical start site (mutants *1_GUA_43_wt_* and *1_wt_43_GUA_*), we saw similar or even higher levels of cell-surface NA (Fig. 5D, top left). The slight increase could be because the amino acid mutations in these two constructs could increase NA expression or stabilize the protein (it is not possible to make a synonymous mutation to the downstream start site since AUG is the only codon for methionine). Such an increase in NA expression or stability is consistent with previous observations that a variety of amino acid mutations increase cell-surface NA levels [91–95].

We also quantified the levels of NA’s enzymatic activity. As has been described previously [91], this can be done simply by transfecting cells with NA protein-expression plasmids and then measuring the total cell-surface enzymatic activity using the fluorogenic surrogate substrate MUNANA [96]. The results of the enzymatic activity assay mirror the surface expression (Fig. 5D, top left). Specifically, the mutant with only the downstream start site has ~34% of the cell-surface activity of the wildtype NA, and this activity is largely dependent on having an AUG at position 43.

To confirm these results in another viral strain, we generated a similar set of mutants of the NA from the A/California/04/2009 (CA09) strain, which is a human isolate from the pdmH1N1 lineage. The results for this NA were broadly similar to those for the WSN strain: mutants without the canonical start codon still had detectable cell-surface protein and activity, which were largely dependent on having an AUG codon at position 43 (Fig. 5D, right panels). Therefore, the AUG at position 43 of NA is capable of initiating translation in multiple viral strains.

The results in Fig. 5D demonstrate that that the codon at position 43 can initiate translation in the absence of the canonical start site. However, because the assays used in those experiments cannot disambiguate between full-length NA and NA43, they do not fully validate the ribosome profiling data suggesting that the codon at position 43 initiates translation even in the presence of the canonical start site. We therefore performed Western blots to directly distinguish between full-length NA and NA43 on the basis of the size of the resulting protein.

NA43 is only 14 amino acids shorter than the 453 amino acid full-length NA. This size difference is difficult to resolve by Western blotting, so we created synthetic constructs by fusing the 5′ end of the NA gene (the 5′ UTR and the first 120 nucleotides of NA) to a short histone (H2B) coding sequence and a C-terminal V5 tag which results in a 180 amino acid reporter. Following the stop codon of the V5 tag, we retained the rest of the NA coding sequence. We again created variants of these synthetic constructs in which the individual AUGs were mutated to codons that are not near-cognate starts. Upon transfecting these constructs into HEK293T cells, the wildtype NA construct shows two bands (lane 2 of Fig. 5E) which run at the expected sizes for proteins initiated at the canonical AUG and the downstream AUG43. Constructs that lack AUG43 only show the band for full-length NA (lanes 3 and 4 of Fig. 5E), and a construct that lacks the canonical AUG start site only shows the band for NA43 (lane 5 of Fig. 5E). The mutants lacking an AUG at both the canonical start site and position 43 had an unexpected band intermediate in size between the full length NA and NA43 (lanes 7 and 8 of Fig. 5E). However, mutating a near cognate putative start codon at position 28 (lane 7 of Fig. S16A) largely eliminates this cryptic band.

As an additional check for initiation at AUG43 in the presence of upstream starts, we also generated constructs that introduce a frameshift just preceding position 43, such that any initiation that occurs 5′ to the frameshift will generate products that are not detected by our V5 antibody. Only the construct that contains an AUG at position 43 (and not GUA at position 43) produces a NA43 band (lane 5 versus lane 6 of Fig. S16B). Therefore, we conclude that the AUG at position 43 can initiate translation in both the presence and absence of the canonical AUG start codon.

We next examined the extent to which NA43 could complement NA during viral replication. To do this, we used reverse genetics [97] to attempt to generate WSN influenza viruses with NAs that lacked one or both of the start sites. By far the highest viral titers were obtained by using the wildtype NA (Fig. 5F). However, we also obtained low but detectable viral titers when using NA that lacked the canonical start codon but had the AUG at position 43, but obtained no detectable viral titers when both start codons were mutated (Fig. 5F). The much lower viral titers when using the mutant that just produces NA43 could be due to a combination of reduced expression and the possible failure of NA protein lacking the cytoplasmic tail and a portion of the transmembrane domain to effectively localize to regions of viral budding [98–103]. Notably, there was no detectable change in viral titer if we mutated the AUG at position 43 but maintained the canonical start codon. Therefore, although NA43 can weakly complement for full-length NA, it is not important for viral growth in the presence of full-length NA in cell culture.

The results in Fig. 5F suggest that NA43 does not substantially contribute to viral growth in cell culture, but simply titering viral supernatants generated by reverse genetics is a relatively insensitive way to quantify how mutations affect growth. We therefore performed competition assays between viruses with wildtype NA and NA lacking the start site at position 43, since such assays are a more sensitive way to detect small differences in viral fitness [104]. We performed these assays by mixing virus with wildtype NA with either the *1_wt_43_GUA_* or *1_wt_43_UUA_* mutant in roughly equal proportion, and then allowing the viruses to compete in low-MOI infections in cell culture. We then used deep sequencing to quantify the relative ratio of the wildtype and mutant NAs both prior to extensive secondary replication (10 hours post-infection) or after many rounds of replication (72 hours post-infection). If NA43 is important for viral fitness in cell culture, then we would expect virus with the wildtype NA (which has the start codon for NA43) to become enriched relative to either mutant. However, there was no enrichment of virus with the wildtype NA (Fig. 5G), indicating that NA43 is not detectably important for viral fitness in cell culture in the presence of full-length NA.

We next considered the possibility that NA43 might contribute to viral fitness only *in vivo*. For instance, it is known that NA has additional roles that impact fitness *in vivo*, such as helping the virus access cells in the presence of mucins [105]. We therefore performed similar competition assays in mice, sequencing the viral population in the mouse lung 4 days after a low-dose infection with a mix of the wildtype and mutant viruses. In the mice, we did see enrichment of the wildtype virus relative to the *1_wt_43_GUA_* NA mutant (Fig. 5H), but this was not statistically significant (Fig. 5H legend). Furthermore, the enrichment was present only for the *1_wt_43_GUA_* mutant but not for the *1_wt_43_UUA_* mutant, making it hard to disambiguate effects due to the lack of NA43 from the effects of introducing an amino acid substitution in NA that eliminates the downstream AUG. Therefore, our experiments did not reveal an unambiguous contribution of NA43 to viral fitness in the presence of full length NA either in cell culture or in a mouse model of infection.

### Translation initiation on host transcripts during influenza infection and anti-viral response

Influenza virus infection and the host anti-viral response alter the expression of many host genes. For example, transcript levels of oxidative phosphorylation genes are refractory to host shutoff during influenza infection [27], and several hundred interferon-stimulated genes are induced during the host anti-viral response [106]. We sought to use our Ribo-seq and Ribo-seq + LTM data to examine translation initiation on host transcripts during influenza infection and the host anti-viral response.

Our strategy for calling translation initiation sites (TIS) on host transcripts was similar to the one we used for influenza transcripts (Fig. 3A). We used less stringent criteria to account for the lower read coverage of host transcripts compared to influenza transcripts (see Methods). We validated our TIS calling strategy for host transcripts using data from a previous study [31]. Our calling method resulted in slightly higher number of annotated TIS and lower number of downstream and upstream TIS than in the TISdb database [83] created using the same data (Fig. 6A). This observation is expected from our conservative strategy of using Ribo-seq data in addition to Ribo-seq + LTM data to decrease false positives resulting from library preparation biases.

**Fig 6.**
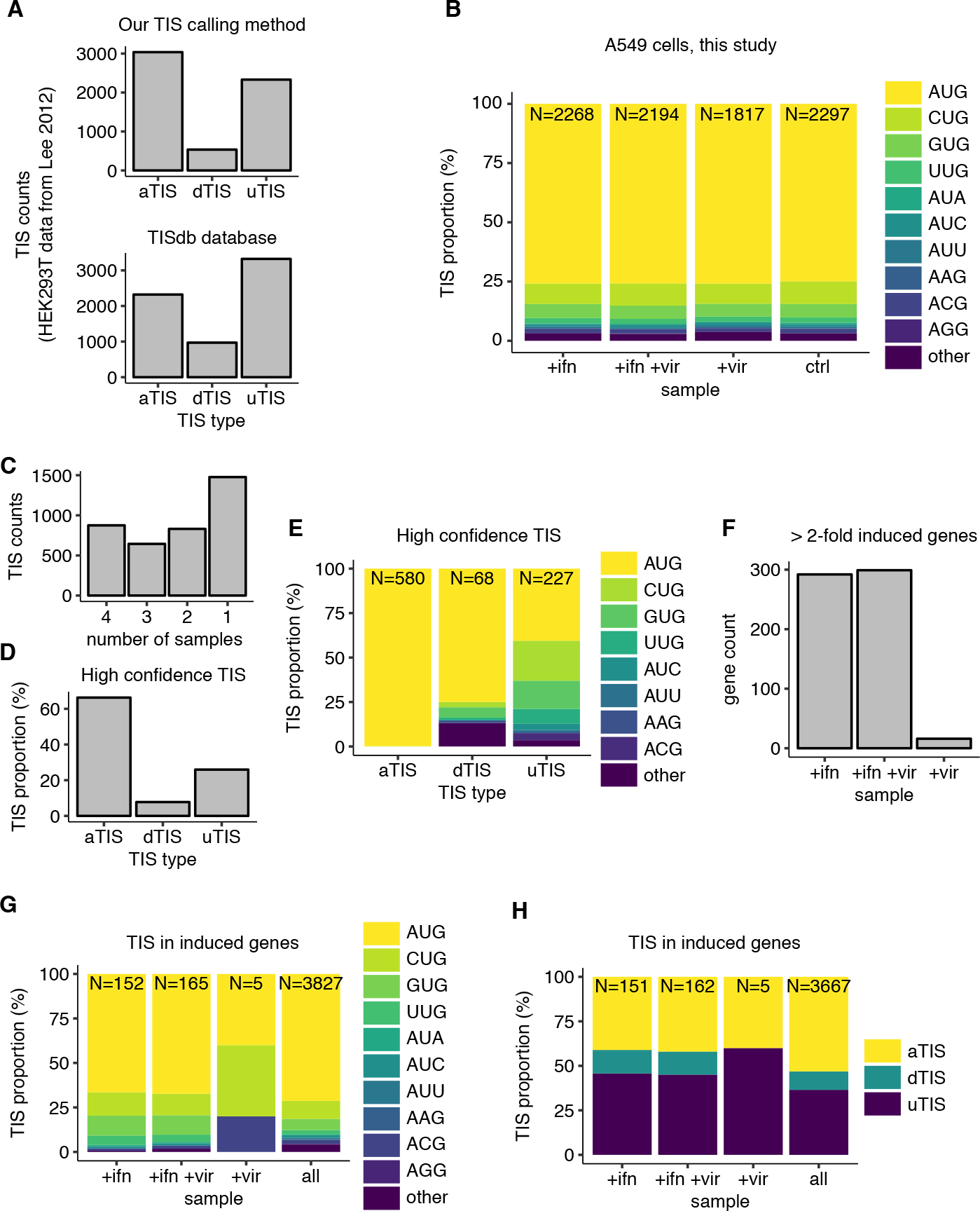
Translation initiation on human transcripts during influenza infection and anti-viral response. **(A)** Number of different TIS types—aTIS, dTIS and uTIS—called using our start site calling method (upper panel) or in the TISdb database [83] (lower panel). See main text for description of our calling method. **(B)** Proportion of different near-cognate AUG codons (or other codons) at the called TIS in each of the four samples in our study. *N* at the top of each bar indicates the total number of TIS called in each sample. **(C)** Overlap among TIS called in each of the 4 samples. Vertical axis indicates the number of TIS that are called in a given number of samples as indicated along the horizontal axis. **(D)** Proportion of each TIS type that are called in all 4 samples (designated as high-confidence TIS). **(E)** Proportion of different near-cognate AUG codons (or other codons) among the high-confidence TIS, stratified by TIS type. *N* at the top of each bar indicates the total number of high-confidence TIS of each type. **(F)** Number of genes that are induced greater than 2-fold (average of RNA-Seq and Ribo-seq counts, median-normalized) upon +ifn, +ifn +vir, or +vir treatment with respect to the control untreated sample. **(G)** Proportion of different near-cognate AUG codons (or other codons) among induced genes shown in F. The set of all TIS called in at least one of the four samples in B is shown as a control. *N* at the top of each bar indicates the total number of TIS that are called in each gene set. **(H)** Proportion of different TIS types among induced genes shown in F. The set of all TIS called in at least one of the four samples in B is shown as a control. *N* at the top of each bar indicates the total number of TIS that are called in each gene set.

Application of our TIS calling strategy to the four samples in our study identified around 2000 candidate TIS in each sample (Fig. 6B). Across all four samples, over 75% of the called TIS were within 1 nucleotide of AUG codons, with CUG and GUG being the next most abundant codons at the called TIS (Fig. 6B). This proportion of AUG and near-cognate AUG codons was similar to those observed with data from HEK293T cells (Fig S17A) [31]. 875 of the called TIS were shared between all 4 of our samples (Fig. 6C), which we designate as high-confidence TIS for further analyses. Among these high-confidence TIS, over 60% corresponded to annotated canonical start codons (Fig. 6D), further validating our start site calling strategy. Less than half of the high confidence TIS identified in our study were shared with those identified from earlier work on HEK293T cells (Fig S17B). This modest overlap in called TIS likely reflects the distinct gene expression landscape between the A549 cells used in our study and HEK293T cells (Fig S17C). The uTIS were highly enriched for near-cognate AUG codons in comparison to dTIS among the high-confidence TIS (Fig. 6E), recapitulating previous observations (Fig. S17D) [30,31].

Previous reporter studies found that non-AUG initiation could be specifically up-regulated under conditions of inflammation or infection leading to antigenic presentation of cryptic peptides [6]. We first sought to test the generality of this observation at the genome-wide level using our called TIS set. Comparison of number of called TIS between our four samples (Fig. 6B) did not reveal a globally higher proportion of non-AUG TIS upon influenza infection, interferon-*β* stimulation, or under both stimuli relative to the uninfected control sample (*P* > 0.05 for all three treatments, binomial proportion test). Since non-AUG TIS tend to be enriched among uTIS and dTIS in comparison to aTIS (Fig. 6E), consistently, we also did not find a higher proportion of uTIS or dTIS upon any of the stimuli relative to the uninfected control sample (*P* > 0. 05 for all three treatments for both uTIS and dTIS, binomial proportion test; Fig. S17E).

We then considered the possibility that the degree of increased non-AUG initiation during influenza infection or interferon-*β* stimulation might be too weak to be detected as a globally higher number of non-AUG start codons in our TIS set called using the Ribo-seq + LTM data. However, even a slightly higher degree of non-AUG initiation during these inflammatory stimuli might favor evolutionary selection for a higher proportion of non-AUG TIS in genes that are specifically up-regulated under these conditions. To test if there is a higher proportion of non-AUG TIS among induced genes, we first identified genes that were induced greater than 2-fold (average of Ribo-seq and RNA-Seq counts) under each of the three treatments in our study. interferon-*β* treatment or interferon-*β* treatment followed by influenza infection resulted in up-regulation of around 300 genes (>2-fold, Fig. 6F). As expected, the most highly induced genes are well-characterized interferon-stimulated genes such as MxA or IFIT1. The genes induced upon either interferon-*β* treatment or interferon-*β* treatment followed by influenza infection show a high degree (> 90%) of overlap (Fig S17F). By contrast, influenza infection on its own resulted in up-regulation of a small set of 16 genes (>2-fold, Fig. 6F) including only a few interferon-stimulated genes. This observation is consistent with recent work showing that activation of immune pathways by influenza virus is rare at the single cell level during infections with viruses that have relatively few defective particles [61].

We examined the TIS codon identity and type in induced genes, and compared it to a control set of all TIS that were called in any of our samples. This analysis revealed that genes that were induced upon either interferon-*β* treatment or interferon-*β* treatment followed by influenza infection had a higher proportion of near-cognate AUG TIS relative to AUG TIS [34% (+ifn induced genes) / 33% (+ifn +vir induced genes) vs 26% (all genes), *P* = 0.02, binomial proportion test, Fig. 6G]. uTIS and dTIS were also present in higher proportion than aTIS among the interferon-induced genes, as expected from the over-representation of non-AUG codons among these TIS types [59% (+ifn induced genes) / 58% (+ifn +vir induced genes) vs 47% (all genes), P < 0.01, binomial proportion test, Fig. 6H]. We did not find the proportion of near-cognate AUG TIS or uTIS and dTIS in the influenza-induced genes to be significantly different from those in all genes (*P* > 0.05, binomial proportion test, Fig. 6G). This lack of statistical significance is likely due to the meager number of called TIS (N=5) in the set of influenza-induced genes. One important caveat that could bias these observations which are based off our Ribo-seq + LTM data is the runoff of elongating ribosomes caused by the lactidomycin treatment. The increased proportion of free ribosomal subunits under these conditions could lead to artifactual detection of initiating ribosomes at start sites that are not normally used in the absence of lactidomycin treatment. However, there is no a priori reason why such artifactual detection will favor an increased proportion of non-AUG initiation specifically on interferon-*β* induced transcripts over the remaining transcripts. Thus our observations are consistent with the hypothesis [6] that near-cognate AUG initiation at non-canonical start sites might be more common in an anti-viral cellular state induced by interferon-*β* treatment. However, in our data this increase arises from the presence of more non-canonical start sites in transcriptionally induced genes, rather than increased non-canonical initiation on existing transcripts.

## Discussion

We have performed the first comprehensive characterization of translation initiation sites on viral and host transcripts during influenza virus infection. In viral transcripts, we identified a total of 14 high confidence translation initiation sites, including 7 of 8 canonical translation initiation sites and two previously characterized non-canonical translation initiation sites: PB1-F2 [14] and a start site at nucleotide 113 of M [21]. The seven alternate start sites that we identified were distributed across the PB1, M, NP, NA, and PA segments, and we found candidate novel N-terminal truncations in NP, PA, and NA.

We biochemically validated one of the new viral alternate start sites, a downstream AUG at codon 43 of NA. We showed that the NA43 protein initiated by this start site is produced even in the context of the canonical start site, is expressed on the cell surface, and is enzymatically active. In addition, this NA43 protein is capable of supporting low levels of viral growth even in the absence of full length NA. The AUG codon at 43 is conserved in most N1 viral lineages. However, we were unable to find any evidence that NA43 impacts viral fitness in cell culture or mice, at least at the relatively low resolution with which viral fitness can be measured in laboratory settings. The N-terminal cytoplasmic tail of NA helps localize the protein to areas of viral budding on the cell surface [98–103]. Therefore, the N-terminally truncated NA43 could localize to different regions of the cell surface than full-length NA. It is interesting to speculate whether such altered localization of the NA43 protein could have some phenotypic significance, such as reducing viral coinfection [107].

One of the motivations for our study was to search for evidence of alternate translation initiation at CUG codons in influenza virus. We hypothesized that initiation at such codons might lead to immune recognition of influenza virus, since CUG codons can initiate immune epitopes [8–13], and initiation at CUG codons is thought to be upregulated during conditions of cellular stress including viral infection [5–7]. We found evolutionary signatures in the viral genome that were consistent with selection against CUG-mediated translation initiation in lineages of influenza that have adapted to mammalian hosts. However, we found minimal experiment support for CUG initiation on influenza transcripts in our ribosome profiling experiments. One possibility is that CUG initiation does generate evolutionarily relevant immune epitopes, but that the levels of CUG initiation are too low to be detected by our experimental design. For instance, CUG initiation could be partially refractory to capture by the translation initiation inhibitor lactidomycin used in our experiments [5], or perhaps only occurs in certain types of cells. In addition, even extremely low levels of translation initiation that are difficult to detect by most experimental methods can generate peptides that can still be recognized by T-cells [25]. A second possibility is that the selection against CUG codons is due to some pressure unrelated to translation initiation. There are other evolutionary signatures of influenza adaptation to mammalian hosts with uncertain origin, such as the decrease in GC content of influenza genome during viral adaptation to mammalian hosts [42].

We also did not find significant shifts in global start site usage towards non-AUG translation initiation during either viral infection of interferon treatment, despite the fact that CUG initiation has been shown to be enhanced in these conditions using biochemical and immunological assays [6]. However, the subset of transcriptionally induced genes upon interferon-*β* treatment did have a significantly higher number of non-AUG translation initiation sites than other genes. Whether these alternate start sites serve any biological function will require further study. One interesting possibility is that the peptides generated from these non-AUG start sites on anti-viral genes could harbor T-cell epitopes that are relevant during the host immune response.

Our ribosome profiling experiments used the translation initiation inhibitor lactidomycin to identify candidate start sites in influenza and host transcripts. One potential concern is that the extended lactidomycin arrest of initiating ribosomes could cause a traffic jam of scanning pre-initiation complexes [108] and lead to promiscuous initiation. Indeed more stringent initiation profiling methods have been developed to address this concern [81]. However, our goal was to detect start sites that might have very low levels of initiation. Hence we focused on achieving the highest possible sensitivity in our assay at the cost of possibly inflating the number of upstream start sites. Further, all the novel start sites identified on influenza transcripts occur downstream of the canonical initiation codon, indicating that traffic jams of pre-initiation complexes is not a major problem in analysis of the viral data. If anything, downstream start sites might be more prone to being missed by our method due to blocks caused by the initiating ribosome at the canonical start codon.

Overall, our work provides the first comprehensive analysis of translation initiation during influenza virus infection. We identified several new alternate translation initiation sites, one of which produces a functionally active viral protein that has not been previously described. However, we found little evidence for large-scale initiation of translation at non-canonical start codons such as CUGs on viral transcripts, and only a modest increase in the proportion of non-canonical and non-AUG start codons on host transcripts upregulated during the anti-viral response. The relative paucity of evidence for virally induced alternate translation initiation in our ribosome profiling experiments is seemingly at odds with many studies [5–13] highlighting its potential role in generating immune epitopes during viral infection, as well as our own evolutionary analyses suggesting evolutionary selection against putative CUG start sites in the viral genome. Further work will be needed to resolve this apparent conundrum and define the extent and significance of alternate translation initiation during viral infection and the resulting immune response.

## Materials and methods

### Data and code availability

All high-throughput sequencing data is available from GEO under accession: GSE114636. Scripts for performing all analyses and generating figures in this manuscript are available at https://github.com/rasilab/machkovech_2018.

### Evolutionary analyses of selection against initiation at CUG codons in influenza

#### Influenza sequence sets

To examine signatures of selection against translation initiation at CUG codons in influenza, we first assembled five influenza sequence sets: human, classical swine, duck, chicken, and human H5N1 influenza. For these sequence sets we only include the protein coding sequences from segments that do not reassort in the human influenza lineage descended from the 1918 lineage: PB2, PA, NP, NS1, M1, M2, and NS2. We retained sequences for a strain if that strain had sequences for all genes in the gene set. If there were multiple sequences for a given strain, we kept just one.

All human, H5N1, swine and avian protein coding sequences were downloaded from the Influenza Virus Resource.
(http://www.ncbi.nlm.nih.gov/genomes/FLU/FLU.html)

For human influenza, we kept coding sequences descended from the 1918 virus, which includes H1N1 from 1918 to 1957, H2N2 from 1957 to 1968, and H3N2 from 1968 to 2011 [109]. Viruses from the 2009 swine-origin H1N1 pandemic were excluded. We subsampled our sequences so that we kept at most 5 randomly chosen strains per year.

For human H5N1 influenza, we obtained sequences from 1997 to 2011, and did not subsample our sequences due to the low number of sequences.

For duck influenza, we obtained sequences from 1956 to 2011. We subsampled our duck influenza sequences so that we kept at most 5 randomly chosen strains per year.

For chicken influenza, we obtained sequences from 1934 to 2011. We subsampled our chicken sequences so that we kept at most 5 randomly chosen strains per year.

For swine influenza we selected classical swine H1N1 influenza viruses isolated in North America between 1934 and 1998.

For alignment and subsequent analysis, we appended the seven protein coding regions (PB2, PA, NP, NS1, M1, M2, and NS2) together for each strain. Sequence sets are aligned to A/Brevig Mission/1/1918 virus using MUSCLE [110], with gaps relative to the A/Brevig Mission/1/1918 strain stripped away. Alignments are provided as S1 File.

#### Number of sequence motifs and motif odds ratio

In Fig. 1A, Fig. 1B, Fig. S1, and Fig. S2B we counted the number of the indicated motif (eg. CUG codons) in the indicated reading frame for PB2, PA, NP, NS1, M1, M2, and NS2 protein coding sequences for the viral strains contained in the human, duck, chicken, swine, H5N1 sequence sets.

We also calculated the motif odds ratio (OR) as in [42]. The OR accounts for the individual nucleotide content of a sequence, and therefore removes the effect of any underlying changes in nucleotide content. The OR is defined as follows (shown for the codon CUG):

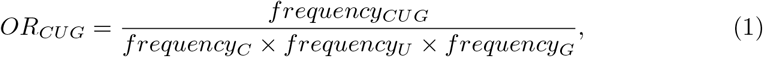

The OR is defined as the frequency of a given motif (CUG) in a sequence divided by the product of the frequency of each nucleotide that comprise the motif (C, U, and G) the sequence. Frequency of a motif such as CUG is defined as the number of codons in a sequence that are CUG divided by the total number of codons in a sequence. Frequency of a nucleotide is the number of a given nucleotide divided by the total number of nucleotides in a sequence. A value greater than 1 indicates there is an excess of the motif given the nucleotide content, and a value less than 1 indicates that there are fewer of the motif given the nucleotide content.

In Fig. S2A, we calculated the odds ratio for CUG for the indicated reading frame in each set of protein-coding sequences for the human, duck, chicken, swine, H5N1 sequence sets.

#### Length of CUG/CUH putative ORFS

For the non-canonical reading frames 1 and 2 of influenza we also examined selection against putative CUG initiation by examining whether there was a host-specific difference in putative ORF lengths (Fig. 1C). We selected a timeframe where the human and swine influenza strains should be reasonably host adapted (1970 to 2011), and took sequences for the human, duck, chicken, swine, H5N1 sequence sets from that time. We calculated the length of all putative ORFs initiated by CUG or CUH (where H is A, U or C) in reading frames 1 and 2 for each influenza sequence set. To plot the data, we took all of the ORF lengths and divided them into 5 ORF length bins. For each influenza host sequence set, we calculated the average number of ORFs for that bin by dividing the number of ORFs in that bin by the number of influenza strains for that host.

#### Consensus motif for CUG start sites

For Fig. S3, we generated a logoplot of the consensus context of CUG start sites called in the ribosome profiling data of Ingolia et al (Supplemental File 3 of [30]).

### Plasmids

#### Influenza reverse genetics plasmids used to generate viral stocks for ribosome profiling

The virus used in the ribosome profiling experiments is a reassortant of the A/PuertoRico/1934 (PR8) and A/WSN/1933 (WSN) influenza strains. NA, HA, M, and NS gene segments are from the WSN strain, encoded by the bidirectional pHW plasmids [97]: pHW186-NA, pHW184-HA, pHW187-M, pHW188-NS. PB1, PB2, and PA segments are from the PR8 strain, encoded by the following plasmids: pHW192-PB1, pHW191-PB2, pHW193-PA. The NP segment was either one of two recoded variants of the PR8 NP, encoded by the following plasmids: pHW-lowCUG-PR8-NP or pHW-highCUG-PR8-NP.

To specifically examine putative alternate initiation at the CUG codons that are selected against in reading frame 0 of influenza, we recoded the PR8 NP to contain either few (low CUG PR8 NP) or many CUG codons (high CUG PR8 NP) (sequences are in S4 File). We specifically chose to recode the CUG content of NP because NP contains many CD8 T-cell epitopes [62,63] and CUG initiation can lead to the generation of CD8 T-cell epitopes [5,6]. Furthermore, we chose PR8 NP as the CD8 T-cell epitopes in PR8 NP for murine models of infection are well characterized [62,111,112]. To generate low CUG PR8 NP, we depleted PR8 NP of the most common alternate start codons AUG, CUG, and GUG as much as possible in all reading frames. We did this because we used low CUG PR8 to generate high CUG PR8, and we wanted to begin with a low background of possible alternate initiation sites. We generated high CUG PR8 NP by adding 20 CUG motifs that occur in reading frame 0 of natural influenza NP sequences to low CUG NP. The mutations we introduced to generate low CUG PR8 NP and high CUG PR8 NP are made with the following constraints: changes must be synonymous with regards to reading frame 0, any synonymous codons that are introduced are chosen to be as frequent as possible in natural existing sequences, and codons must exist in at least 100 of the sequences. The sequences we used for this analysis are all full-length influenza A NP coding sequences from the Influenza Virus Resource. Sequences were aligned using MUSCLE [110], and alignments are included in S2 File.

#### NA43 plasmids for MUNANA and NA surface expression

To measure NA surface expression and NA MUNANA activity in Fig. 5D, we generated constructs by placing NA into an HDM plasmid in which expression of the insert is driven by the CMV promoter. Following the CMV promoter, we included the NA viral 5′ UTR, NA coding sequence, a C-terminal V-5 epitope tag (used for surface staining), and an internal ribosomal entry site (IRES) GFP (used for calculating transfection efficiency).

We made the following set of mutagenized constructs for WSN NA:

pHDM-vUTR-WSN-NA_1ATG-43ATG_V5-IRES-GFP,
pHDM-vUTR-WSN-NA_1ATG-43GTA_V5-IRES-GFP,
pHDM-vUTR-WSN-NA_1ATG-43TTA_V5-IRES-GFP,
pHDM-vUTR-WSN-NA_1GTA-43ATG_V5-IRES-GFP,
pHDM-vUTR-WSN-NA_1GTA-43GTA_V5-IRES-GFP, and
pHDM-vUTR-WSN-NA_1GTA-43TTA_V5-IRES-GFP.

Similar constructs were made for California/04/2009 NA (CA09). In these constructs, we mutated an additional in-frame AUG that was present at coding nucleotides 55-57 to UUA. We did this because the downstream AUG may function as a start site in the absence of either of the other AUG codons, and have the potential to complicate interpretation of our assays if the AUG starting at nucleotide 55 generates a functional NA.

For CA09 we made: pHDM-vUTR-CA09-NA_1ATG-43ATG-55TTA_V5-IRES-GFP,

pHDM-vUTR-CA09-NA_1ATG-43GTA-55TTA_V5-IRES-GFP,
pHDM-vUTR-CA09-NA_1GTA-43ATG-55TTA_V5-IRES-GFP,
pHDM-vUTR-CA09-NA_1GTA-43GTA-55TTA_V5-IRES-GFP.

We also made a construct that lacks the V5 tag as a negative control for background V5 staining (pHDM-WSN-NA-IRES-GFP).

#### NA43 plasmids for Western blots

To examine whether there is initiation at site 43 of WSN NA (Fig. 5E), we generated constructs in an HDM plasmid in which expression of the insert is driven by the CMV promoter and the 5′ UTR of WSN NA. The protein coding sequence consists of codons 1-40 of WSN NA fused to Histone-2B, followed by a C-terminal V5 epitope tag (for immunoblot detection). The stop codon is followed by the remainder of NA and IRES GFP. We made the following constructs:

pHDM-vUTR-WSN-NA-aa1-40_1ATG-43ATG_H2B-V5-IRES-GFP,
pHDM-vUTR-WSN-NA-aa1-40_1ATG-43GTA_H2B-V5-IRES-GFP,
pHDM-vUTR-WSN-NA-aa1-40_1ATG-43TTA_H2B-V5-IRES-GFP,
pHDM-vUTR-WSN-NA-aa1-40_1GTA-43ATG_H2B-V5-IRES-GFP,
pHDM-vUTR-WSN-NA-aa1-40_1GTA-43GTA_H2B-V5-IRES-GFP, and
pHDM-vUTR-WSN-NA-aa1-40_1GTA-43TTA_H2B-V5-IRES-GFP.

We made a construct that lacks the 5′ viral UTR and the first 42 coding nucleotides of NA, so begins at the AUG at coding nucleotide 43 which serves as a size control for NA43 (pHDM-WSN-NA-aa14-40-H2B-V5-IRES-GFP). We also made several constructs to eliminate cryptic products that appear in constructs lacking an AUG at both the canonical start and position 43.

pHDM-vUTR-WSN-NA-aa1-40_1GTA-28GTT-43GTA_H2B-V5-IRES-GFP mutates a near cognate start codon at position 28.

pHDM-vUTR-WSN-NA-aa1-40_1GTA-35Tinsert-43GTA_H2B-V5-IRES-GFP and pHDM-vUTR-WSN-NA-aa1-40_1GTA-35Tinsert-43ATG_H2B-V5-IRES-GFP contain a T at position 35 to introduce a frameshift.

#### NA43 plasmids for viral rescue

We used the bidirectional pHW plasmids [97] for all WSN segments (pHW181-PB2, pHW182-PB1, pHW183-PA,pHW184-HA, pHW185-NP, pHW187-M, pHW188-NS) except NA. For NA we used pHH [113], which includes only the Pol I promoter, so that NA mRNA would only be made from the native vRNA. We generated the following set of mutagenized constructs: pHH-WSN-NA_1ATG-43ATG,

pHH-WSN-NA_1ATG-43GTA, pHH-WSN-NA_1ATG-43TTA,
pHH-WSN-NA_1GTA-43ATG, pHH-WSN-NA_1GTA-43GTA,
pHH-WSN-NA_1GTA-43TTA.

#### NP plasmids for Western blots

To examine whether there is initiation at site 322 of NP (Fig. 5E), we generated constructs in an HDM plasmid in which expression of the insert is driven by the CMV promoter. Following the CMV promoter, we include the 5′ UTR of the indicated NP followed by a C-terminal 3x FLAG epitope tag (for immunoblot detection). We made the following constructs: pHDM-vUTR-lowCTP-PR8-NP-FLAG, pHDM-vUTR-highCUG-PR8-NP-FLAG, pHDM-vUTR-wt-PR8-NP-FLAG. We made a construct that lacks the 5′ viral UTR and begins at the AUG at coding nucleotide 322 which serves as a size control for NP322 (pHDM-322start-wt-PR8-FLAG).

### Cells and cell culture media

We used the human lung epithelial carcinoma line A549 (ATCC CCL-185), the human embryonic kidney cell line 293T (ATCC CRL-3216), the MDCK-SIAT1 variant of the Madin Darby canine kidney cell line overexpressing human SIAT1 (Sigma-Aldrich 05071502), and MDCK-SIAT1-TMPRSS2 variant cells which express both SIAT1 and the protease TMPRSS2 [114].

All cell lines were maintained in D10 (DMEM supplemented with 10% heat-inactivated fetal bovine serum, 2 mM L-glutamine, 100 U of penicillin/ml, and 100 *μ*U,g of streptomycin/ml). Experiments were performed with D10 unless otherwise indicated.

WSN growth media contains Opti-MEM (Gibco) supplemented with 0.05% heat inactivated FBS, 0.3% BSA, 100 U of penicillin/ml, 100 *μ*g of streptomycin/ml, and 100 *μ*g of calcium chloride/ml).

### Virus generation by reverse genetics

#### Generation of virus for ribosome profiling

A co-culture of 293T and MDCK-SIAT1 cells seeded at 4 × 10^5^ and 0.5 × 10^5^ cells per well respectively in 6-well dishes were transfected using BioT and a mixture containing 250 ng of each of the eight plasmids encoding WSN NA, HA, M, NS (pHW186-NA, pHW184-HA, pHW187-M, pHW188-NS), PR8 PB1, PB2, PA (pHW192-PB1, pHW191-PB2, pHW193-PA), and either low or high CUG PR8 NP (pHW-lowCUG-PR8-NP or pHW-highCUG-PR8-NP). We refer to these viruses as low CUG NP WSN and high CUG NP WSN. At 20 hours post transfection, we changed the media from D10 to WSN growth media. At 48 hours post-transfection we collected the viral supernatant, centrifuged the sample at 2000*g* for 5 minutes to remove cellular debris, and stored aliquots of the clarified viral supernatant at −80 °C. We titered the virus by TCID50 (using MDCK-SIAT1 cells). Viruses were expanded starting from an MOI of .05 for 55 hr. The resulting expanded virus was collected and stored in the same manner. The expanded virus was titered by HA staining (using A549 cells) and TCID50 (using MDCK-SIAT1) and used for ribosome profiling.

#### Generation of viruses for NA43 viral growth experiments

A co-culture of 293T and MDCK-SIAT1-TMPRSS2 cells seeded at 4 × 10^5^ and 0.5 × 10^5^ cells per well in 6-well dishes were transfected using BioT and a mixture containing 250 ng of each of the eight plasmids encoding WT WSN PB2, PB1, PA, HA, NP, M, NS (pHW181-PB2, pHW182-PB1, pHW183-PA, pHW184-HA, pHW185-NP, pHW187-M, pHW188-NS) and the appropriate NA variant (pHH-WSN-NA_1ATG-43ATG, pHH-WSN-NA_1ATG-43GTA, pHH-WSN-NA_1ATG-43TTA, pHH-WSN-NA_1GTA-43ATG, pHH-WSN-NA_1GTA-43GTA, pHH-WSN-NA_1GTA-43TTA). MDCK-SIAT1-TMPRSS2 cells were used as it enables HA activation in the absence of NA activity, thus allowing us to minimize indirect effects of NA absence. Viruses were rescued in biological triplicate with three separate minipreps of each NA construct. At 20 hours post transfection, we changed the media from D10 to WSN growth media. We collected 500 ul aliquots of virus 48 and 72 hours post transfection. We titered the virus by TCID50 (using MDCK-SIAT1 TMPRSS2 cells). Viruses were expanded starting from an MOI of .01 for 48 hours. We titered the expanded virus by TCID50 (using MDCK-SIAT1-TMPRSS2 cells).

### Virus titering

#### By HA staining

We infected A549 cells in WSN growth media for 10 hours, collected cells, and stained for HA using H7-L19 antibody [115,116] at 10 ug/ml, followed by goat anti-mouse IgG secondary antibody conjugated to APC, and analyzed by flow cytometry to determine the fraction of cells that were HA positive. We used the Poisson equation to calculate viral titer with respect to HA expressing units.

#### By TCID50

We titered virus by TCID50 on MDCK-SIAT1 or MDCK-SIAT1-TMPRSS2 expressing cells and calculated viral titer using the Reed-Muench formula [117] implemented here: https://github.com/jbloomlab/reedmuenchcalculator.

### Ribosome profiling

#### Ribo-seq library generation

We performed ribosome profiling (Ribo-seq and Ribo-seq + LTM) with four different samples: untreated A549 cells, interferon-*β* treated A549 cells, influenza virus infected A549 cells, interferon-*β* pre-treated and influenza infected A549 cells (Fig. 2A). For each sample, we used two 15 cm plates of A549 cells, seeded with 7 × 10^6^ cells 16 hours prior to start of treatment such that cells were 70-80% confluent at harvest. Five hours before influenza infection, the medium in each plate was replaced with fresh D10 growth media either with or without interferon-*β* (at 20 U/ml). At the time of viral infection, the growth media was removed and replaced with four different WSN growth media corresponding to the four different samples in our experiment. For the +vir sample, we used concentrated virus stocks comprising approximately 2% of the growth media added to cells. Since virus was grown on MDCK-SIAT1 cells, we added the same volume of MDCK-SIAT1 supernatant as the viral stock to the uninfected sample (ctrl sample). For the +ifn sample, we used MDCK-SIAT1 supernatant supplemented with 20 U/ml of interferon-*β* (PBL 11415). For the +ifn +vir sample, we mixed the viral supernatant with interferon-*β* and added it to cells.

Influenza infections consisted of a 1:1 mixture of high CUG NP WSN and low CUG NP WSN viruses at a total dosage of 1 HA-expressing units / cell as calculated using A549 cells not treated with interferon-*β*. This corresponded to an MOI of approximately 30 TCID50 per cell as measured using MDCK-SIAT1 cells. These infection conditions resulted in approximately 70% HA high positive cells for the +vir sample. Samples were harvested 5 hours after influenza infection for ribosome profiling library preparation.

Ribosome profiling protocol was adapted from [118] with the following modifications. For our Ribo-seq + LTM samples, we removed the treatment medium from each plate and added media back containing LTM at 5 *μ*M. These LTM-containing plates were then incubated for 30 min to allow runoff of elongating ribosomes [31]. For sample harvesting, we removed media from each plate and flash froze samples by placing the plate in liquid nitrogen and transferred to −80 °C until lysis. We took a portion of the cell lysate from our Ribo-seq samples for RNA-Seq library preparation. For ribosome profiling, we performed nuclease footprinting treatment by adding 600U RNase I (Invitrogen AM2294) to 18 *A*_260_ units of RNA [119]. We collected ribosome protected mRNA fragments using MicroSpin S-400 HR Columns (GE Healthcare 27-5140-01) following instructions from the Art-Seq ribosome profiling kit (RPHMR12126, Illumina), and purified RNA from the flow through for size selection. We gel-purified ribosome protected fragments with length between 26 and 34 nucleotides using RNA oligo size markers. We used polyA tailing instead of linker ligation following previous studies [26,31]. Finally, we prepared RNA-seq libraries using the NEBNext Ultra Directional RNA Library Prep Kit (NEB E7420) after depleting ribosome RNA using the Ribozero Gold kit (Illumina MRZG126). Ribo-seq and RNA-seq libraries were sequenced on an Illumina HiSeq 2500 in 50bp single end mode resulting in between 15-40 million reads per sample (see Fig. S5A).

#### Pre-processing and alignment of sequencing data

For Ribo-seq libraries, we trimmed polyA tails using cutadapt 1.12 [120] with parameters `--adapter=AAAAAAAAAA --length=40 --minimum-length=25` to retain trimmed reads that were between 25 and 40 nucleotides in length. For RNA-seq libraries, we first reverse complemented the read to obtain the sense orientation using `fastx_reverse_complement 0.0.13` (http://hannonlab.cshl.edu/fastx_toolkit/) with parameters `--Q33,` and then trimmed reads to 40 nucleotides. To remove rRNA, we discarded trimmed reads aligning to four human rRNA sequences (28S:NR_003287.2, 18S:NR_003286.2, 5.8S:NR_003285.2, and 5S:NR_023363.1) from the `hg38` genome. The alignment was done using `bowtie 1.1.1` [121] with parameters `--seedlen=23 --threads=8`. We aligned the remaining non-rRNA to human transcripts (Gencode v24) using rsem 1.2.31 [122] with parameters `--output-genome-bam --sort-bam-by-coordinate`. We also aligned the non-rRNA reads to an influenza genome containing both low and high CUG NP using the above rsem parameters including the extra options `-seed-length 21 -bowtie-n 3`. The influenza genome file is provided in S4 File and the corresponding annotations in S5 File.

#### Calculation of influenza and host transcript coverage

The posterior probablilty score from `rsem` (`ZW` field in the `BAM` output) was used to calculate coverage for reads on transcripts. For influenza reads, we considered only reads that map to the positive sense of influenza transcripts for all analyses except where explicitly indicated otherwise. To obtain higher resolution mapping of ribosome protected fragments, we performed variable trimming based on fragment length, reads that were between 25 and 32 nucleotides in length were assigned to a P-site at 14 nucleotides from the 5′ end of the read, and reads between 33 and 39 nucleotides were assigned to a P-site at 15 nucleotides from the 5′ end of the read.

#### Ribo-seq start or stop codon aligned read aggregation plots

For human transcripts, a set of non-redundant protein-coding transcripts was compiled by using the `gencode v24` annotations and applying the following criteria: The corresponding CDS must be part of the Consensus CDS (CCDS) project; It must be labeled as a `principal transcript` by APPRIS; For each gene, we selected the lowest numbered CCDS ID; For each CCDS, we selected the lowest numbered transcript ID.

For aggregation of sequencing reads, we further selected the set of transcripts requiring that each transcript has a minimum coverage of 0.33 reads per nucleotide in the coding region in a given sample. For each ribosome P-site in a transcript, the normalized ribosome density value was calculated by dividing the P-site read count by the average P-site read count across the coding region of that transcript. The normalized P-site density across all transcripts was calculated by averaging each P-site position (aligned relative to the start or stop codon).

#### Proportion of Ribo-seq reads aligning to each reading frame

Normalized P-site density across reading frames was calculated for the coding regions of the non-redundant set of transcripts included in the analysis used for read aggregation plots. The single- and dual-coded regions of M, NS and PB1 were parsed from S5 File, and the normalized P-site density across each of the regions was plotted.

#### Calling and analysis of candidate TIS

We used a zero-truncated negative binomial distribution (ZTNB) to statistically model the background distribution of Ribo-seq and Ribo-seq + LTM counts in transcripts with more than 50 positions with non-zero counts [80,81]. We first added the Ribo-seq and the Ribo-seq + LTM read counts of the two neighboring positions to each position in the genome (referred to as pooled counts below). This was done in order to account for the +/−1 nucleotide uncertainty in the P-site assignment.

Candidate start sites were identified based on the following criteria: For influenza transcripts, the ZTNB-based P-value for the Ribo-seq + LTM pooled counts at that location must be <0.01 and 1000-fold higher than the P-value of the Ribo-seq pooled counts at the same location (Fig. 3A, left panel), or must have an absolute P-value less than 10^−7^. For host transcripts, we required the Ribo-seq + LTM P-value to be only 100-fold higher than the P-value of the Ribo-seq pooled counts. Additionally, for host transcripts, we required that the read counts be greater than an absolute threshold across all transcripts. This threshold was estimated by requiring P<0.05 in a ZTNB model fit to the bottom 99% of all non-zero P-site Ribo-seq + LTM pooled counts across all transcripts. Only the highest pooled counts within each 30 nucleotide window was called as a candidate TIS. From the called TIS, we assigned the identity of the start codon by looking at a window -1 to +1 nucleotides from the TIS peak and assigning the start codon based on following hierarchy: AUG, CUG, GUG, UUG, AUA, AUC, AUU, AAG, ACG, AGG, and other. If there are multiple near cognate codons in the window, the codon was assigned based on the order in the above list.

For Fig. 3H and 3I, we consider the canonical frame defined by the aTIS as frame 0, and we designate any ORF out of frame with the aTIS as an out-of-frame ORF. For candidate host TIS, we designate the start codon of the canonical transcript of each gene (as defined above) as the annotated TIS. uTIS, dTIS, and the frame of their ORF are identified with respect to the annotated TIS. Host genes that have a median-normalized fold-change in average Ribo-seq and RNA-seq counts greater than 2-fold upon +ifn, +vir, or +ifn +vir treatment are considered as induced genes under that condition.

For analysis of data from [31], we downloaded the raw sequencing data from the Sequence Read Archive, BioProject PRJNA171327, and analyzed it using the same pipeline that we used for our data. The same pipeline was also used for analysis of the raw sequencing data from [27] (NCBI Gene Expression Omnibus, GSE82232). The only difference from the analysis of our dataset is that we identified P-site as the 13th nucleotide from the 5′ end of the read for both these datasets.

Binomial proportion test for comparing proportions of different TIS was done using the `R` function `prop.test` with the alternate hypothesis set to `greater`.

#### Examining initiation at CUG codons in high and low CUG NP variants

In Fig. 4A, we considered reads that map with zero mismatches to either low or high CUG NP or both NP variants. We designated reads as belonging to one of the variants if it spans a SNP (recoded CUG codon), and thus could be identified uniquely. Coverage of non-unique, low CUG, and high CUG variants is shown as a stacked bar blot. For Fig. 4B, we plot the ratio of P-site count between the high and low CUG variants at each NP position to the mean value of the same quantity across the two variants. All P-sites have a pseudocount of 1 added to both the low and high CUG read counts.

#### NA surface expression and MUNANA NA activity assay

We performed NA surface expression and NA activity assays (Fig. 5D), as described in [90,91]. Briefly, we transfected 293T cells seeded at 2 × 10^5^ cells per ml in 12-well plates using BioT and 1000 ng of each WSN or CA09 pHDM-vUTR-NA-V5-IRES-GFP construct, including a control pHDM-vUTR-NA-V5-IRES-GFP (lacking a V5 tag), and a no vector control. Unless indicated, we performed assays in triplicate from independent minipreps of constructs. At 20 hour post transfection, we trypsinized, collected, and resuspended cells in a non-lysing buffer - MOPS buffered saline (15 mM MOPS, 145 mM sodium chloride, 2.7 mM potassium chloride, and 4.0 mM calcium chloride, adjusted to pH 7.4 and supplemented with 2% FBS). We divided each sample in half to perform NA surface expression and MUNANA NA activity assays in parallel.

#### NA surface expression

The transfected 293T cells were stained for cell-surface NA using an anti-V5 AF647-conjugated antibody (Invitrogen 45-1098) at a 1:200 dilution. Cells were analyzed by flow cytometry to determine the MFI of AF647 (APC channel) among GFP-positive (transfected) cells. Reported values were normalized to the *1_wt_43_wt_* NA with the background value from the pHDM-vUTR-NA-V5-IRES-GFP (lacking V5 tag) subtracted.

#### MUNANA NA activity assay

The transfected 293T cells were used to assay for NA activity using MUNANA substrate (Sigma M8639). We performed the MUNANA assay in a non-lysing buffer (MOPS buffered saline supplemented with 2% FBS). We used 2 dilutions of our resuspended cell sample (25% and 2.5% of original total sample volume) to ensure that we are within the dynamic range of the assay. We performed the assay in black 96-well plates in a 200 ul volume. We added MUNANA to a concentration of 150 *μ*M and incubated samples for 45 min at 37 “C, and quenched by adding 50 *μ*l of 142 mM NaOH in 100% ethanol. Fluorescence was measured at an excitation wavelength of 360 nm and an emission wavelength of 448 nm. Activity was calculated by subtracting the background value of untransfected cells, correcting for transfection efficiency (% GFP positive from NA surface expression assay), and normalizing to *1_wt_43_wt_* NA.

### Western blots

We transfected 293T cells seeded at 2 × 10^5^ cells/well in a 12-well plate with the pHDM-vUTR-WSN-NA-aa1-40-H2B-V5-IRES-GFP constructs or pHDM-vUTR-NP-FLAG constructs. We collected cell lysates 20 hour post transfection, using RIPA buffer containing (1% NP40, 1% Sodium deoxycholate, 0.1% SDS, 150 mM NaCl, 50 mM Tris (pH 8), 0.1 mM EDTA, and Roche cOmplete ULTRA Tablet Protease Inhibitor).

For NA43, lysates were run on a 16.5% tris-tricine gel. For NP, lysates were run on a 4-20% tris-glycine gel. NA was detected using 1:2500 dilution of anti-V5 antibody (Invitrogen R960), followed by a 1:2500 dilution of DyLight 800 Rabbit anti-Mouse (Invitrogen SA5-10164). NP was detected using 1:2500 dilution of anti-FLAG antibody (Sigma F1804), followed by a 1:2500 dilution goat anti-mouse Alexa-Fluor 680 (Invitrogen A-21058). For loading controls, we used either GAPDH or H3. We used anti-GAPDH at a 1:2500 dilution (RD systems AF5718), followed by a 1:2500 Alexa-Fluor 680 donkey anti-goat secondary (Invitrogen A-21084). We used anti-H3 at a 1:10000 dilution (abcam 1791), followed by a 1:10000 Alexa-Fluor 680 goat anti-rabbit secondary (Invitrogen A-21109). Western blots were imaged using the LI-COR imaging system.

### NA43 codon analysis

To examine the conservation of the AUG at coding site 43 in Fig. 5C, we downloaded all full-length N1 NA protein-coding sequences from the Influenza Virus Resource. We used phydms [123] to construct a codon-level alignment in reference to the WSN sequence used in the ribosome profiling experiment. The alignment is subsampled such that all sequences have at least 2 amino acid differences relative to other sequences. This is to avoid having the alignment dominated by many highly similar sequences that are heavily represented in the database. We parse sequences into the following four sets: human seasonal H1N1 sequences isolated before 2009, human pandemic H1N1 sequences isolated after 2009, all avian sequences, and all swine sequences. We further subset human seasonal H1N1 sequences so that we only keep 1 sequence per year. Sequences are in S3 File. We then count the number of each codon at coding nucleotides 43-45 for our set of sequences. We consider CUG, GUG, UUG, ACG, AGG, AUC, AUU, AAG, AUA as near cognate start codons.

### Competition assays to examine impact of NA43 on viral fitness

For the competition assays (Fig. 5G and H) to examine if NA43 conferred a viral fitness advantage, we made 6 mastermixes of virus containing 1:1 mixes (as measured by TCID50) containing either *1_wt_43_wt_* and *1_wt_43_GUA_* (N=3) or *1_wt_43_wt_* and *1_wt_43_UUA_* (N=3) virus. For each virus, we used 3 biological replicates of expanded *1_wt_43_wt_*, *1_wt_43_GUA_* and *1_wt_43_UUA_* virus. The same viral mastermixes were used for both cell culture and mouse competition assays The viral mastermixes underwent 1 freeze thaw cycle prior to the mouse experiment.

#### Cell-culture NA43 competition assays

Competitions were done at a low total MOI of 0.001 (equivalent to an MOI of 0.0005 for each individual viral variant) in MDCK-SIAT1-TMPRSS2 cells. For each competition pair, we used one well of a 6-well plate (seeded at 5 × 10^5^cells per well) and one 10 cm dish (seeded at 1.4 × 10^6^cells per plate). We harvested the 6-well plate for cellular RNA at 10 hour post infection to get an early estimate of the viral ratios. We let the competitions proceed for 72 hours, and at 72 hours we collected 700 *μ*l of virus supernatant from the 10 cm dish. Competitions were performed in WSN growth media. For the 10 hour timepoint, we extracted RNA using the Qiagen RNeasy mini plus kit, following the manufacturer’s protocol. For the viral supernatant samples from the cell culture competition, we extracted viral RNA using the Qiagen QIAamp Viral RNA Mini Kit, following the manufacturer’s protocol (using 140 *μ*l virus containing supernatant).

#### Mouse NA43 competition assays

All animal experiments were conducted in accordance with National Institutes of Health guidelines for humane treatment of animals and were approved by the Institutional Animal Care and Use Committee of the Fred Hutchinson Cancer Research Center.

For each viral mastermix, we innoculated 3 female BALB/c mice (technical replicates) with 20 *μ*l of virus mastermix (containing 2000 TCID50). BALB/c mice were 6-8 weeks old, from Jackson labs. For infections, mice were first anesthetized with 0. 2 mg ketamine and 20 *μ*g xylazine. Mice were weighed daily and lungs were harvested and flash frozen on day 4 post infection.

We homogenized whole lung in 2.4 ml buffer RLT using the gentleMACS dissociator. The homogenate was clarified by centrifugation, and 700 *μ*l of supernatant was used for RNA extraction using the Qiagen RNeasy kit, following the manufacturer’s protocol.

#### Sequencing to determine NA43 mutant frequency

We reverse transcribed NA vRNA gene from the extracted RNA using SuperScript III Reverse Transcriptase for first-strand cDNA synthesis. The reverse transcriptase reaction contained 500 ng cellular RNA template or 4 *μ*l viral RNA and used the primer U12-NA-F (5-AGCGAAAGCAGGAGTUUAA-3) at 5 *μ*M. We then carried out targeted deep sequencing of the mutated region in NA. We first amplified a region of the NA gene that encompassed position 43 using 1.5 *μ*l cDNA template and the primers U12-NA-F and NA-249-R (5-GGACAAAGAGAUGAATTGCCGG-3). We used the following PCR program, 95 °C 2 min, followed by 27 cycles of: 95 °C 20s, 70 °C 1s, 50 °C 30s, 70 °C 20s.

We purified the PCR product using 1.5X AMPure XP beads (Beckman Coulter) We performed a second round of PCR to add a portion of the Illumina adapters using 3 ng DNA and the following PCR primers: Rnd1-F-WSN-NA (5-CTTTCCCTACACGACG CTCTTCCGATCTAGCGAAAGCAGGAGTUUAA-3) and Rnd1-R-WSN-NA (5-GGA GTTCAGACGTGTGCTCTTCCGATCUGGACAAAGAGAUGAATTGCCGG-3).
We used the following PCR program: 95 °C 2 min, followed by 8 cycles of: 95 °C 20s, 70 °C 1s, 54 °C 20s, 70 °C 20s.

We purified the PCR product using 1.5X AMPure XP beads and used 1.5 ng of this product as template for a third round of PCR using the following pair of primers that added the remaining part of the Illumina sequencing adaptors as well as a 6 or 8-mer sample barcode (xxxxxx): (5-AAUGATACGGCGACCACCGAGATCTACACTCTTT CCCTACACGACGCTCTTCCGATCT-3) and Rnd2-R (5-CAAGCAGAAGACGGCA TACGAGATxxxxxxGTGACUGGAGTTCAGACGTGTGCTCTTCCGATCT-3). We used the following PCR program: 95 “C 2 min, followed by 6 cycles of: 95 °C 20s, 70 °C 1s, 58 °C 10s, 70 °C 20s.

The resulting DNA libraries were sequenced on an Illumina Hi-Seq. We processed the sequencing reads to determine the frequency of the wildtype NA and mutant NA in each competition as follows. To determine the frequency of wildtype NA, we aligned our samples to the *1_wt_43_wt_* sequence, and counted the number of reads containing AUG at coding nucleotides 43-45. To determine the frequency of mutant NA, we aligned our samples to etiher the *1_wt_43_GUA_* or *1_wt_43_UUA_* sequence, and counted the number of reads containing GUA or UUA at coding nucleotides 43-45. We used `dms_tools` [124] for alignment of sequence data, and allowed only 2 nucleotide mismatches over coding nucleotides 1-48.

#### Analysis of NA43 mutant frequency

To examine if there is an enrichment of wildtype viruses mutant over time, we first calculated the ratio of wildtype verses mutant at each timepoint using the counts of AUG, UUA, or GUA codons at postion 43-45. We then calculated the enrichment of wildtype relative to mutant over time. To do this we divided the ratio of wildtype to mutant at the endpoint (either 72 hour cell-culture or mouse value) by the ratio of wildtype to mutant at the 10 hour cell-culture timepoint. We compared the endpoint to the 10 hour timepoint instead of assuming that the initial ratio is 50:50 as the precision of TCID50 is less than that of deep sequencing, and the 10 hour timepoint allows us to measure relative levels of infectious virus for each variant. A value greater than 1 indicates an enrichment of wildtype over time. To assess significance, we performed two one-sided paired t-tests (mouse *1_wt_43_GUA_* and mouse *1_wt_43_UUA_*). For the t-test, we used the log transformed ratios of wildtype to mutant at each timepoint. For the mouse assay, we first took the mean of the 3 technical replicates (using the log transformed value), and paired the average mouse value for each biological replicate to the 10 hour cell culture timepoint. The sequencing counts of codons and the resulting ratios used in analysis are included in S1 Table.

## Acknowledgments

We thank Adam Geballe, Ji Wan, & Shu-Bing Qian for helpful discussions. The work of ARS is supported by grant R35 GM119835 of the NIGMS of the NIH. The work of JDB is supported by grant R01 AI127893 of the NIAID of the NIH, and a Faculty Scholars Award from HHMI and the Simons Foundation. HMM was supported in part by grant T32 AI083203 of the NIAID of the NIH.

## Supporting information

**Fig S1.**
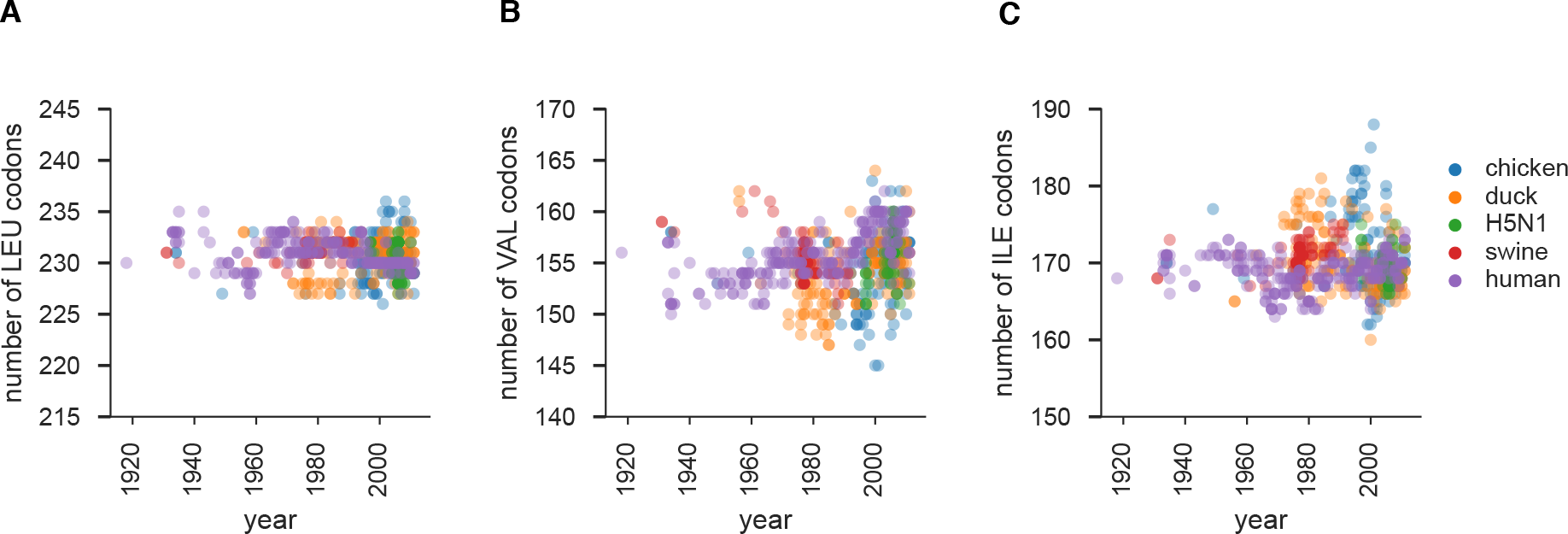
Evolution of amino acid usage in influenza. The number of LEU, VAL, ILE codons in reading frames 0 of the influenza genome over time in human, avian, and swine lineages. There is no systematic trend for enrichment or depletion of any of the amino acids.

**Fig S2.**
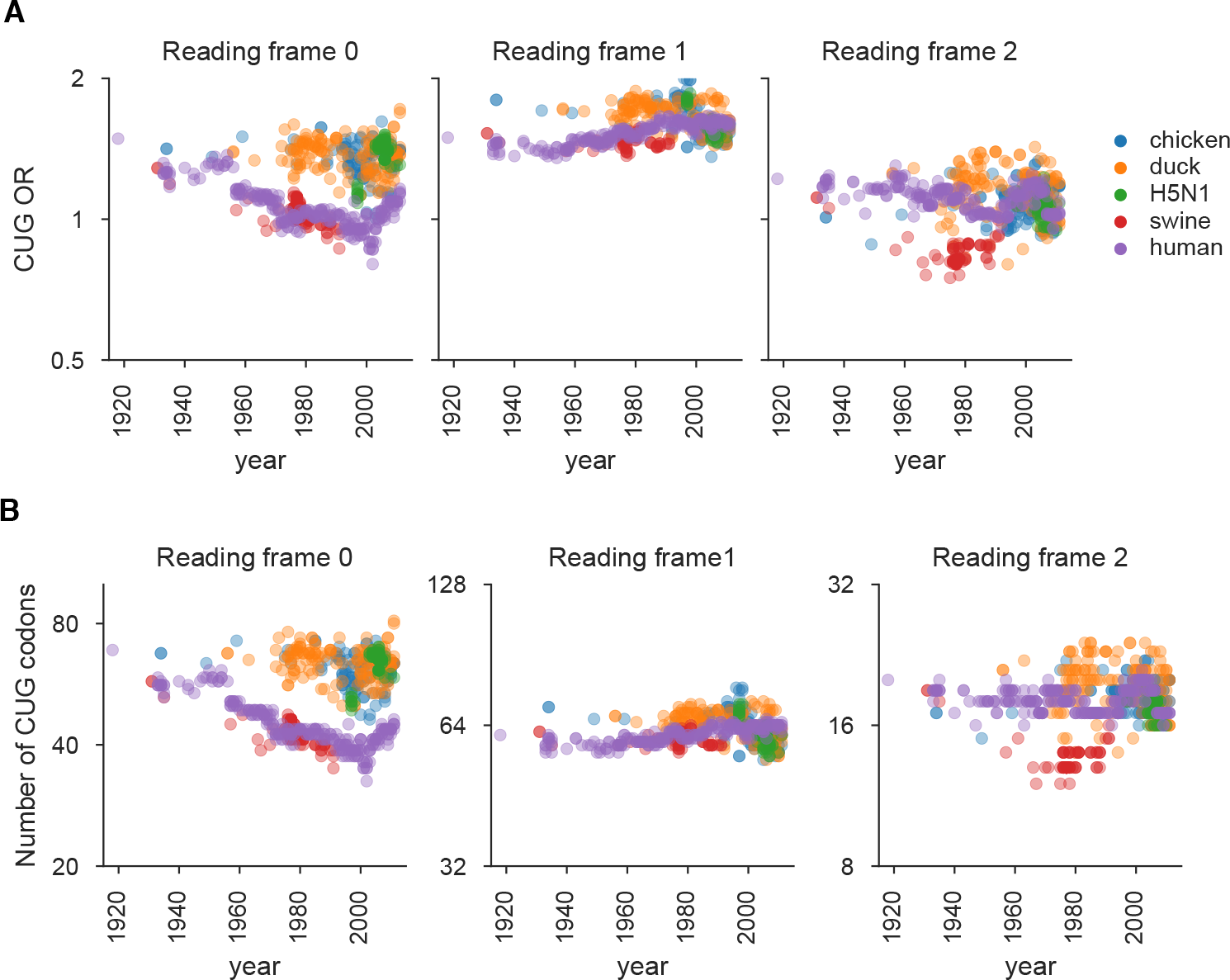
Evolution of CUG codons in influenza. **(A)** Evolution of the CUG odds ratio 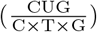 in reading frames 0, 1, and 2 over time in human, avian, and swine lineages. The selection against CUG in reading frame 0 exists even when we use the odds ratio to correct for nucleotide usage. **(B)** Evolution of the number of CUG codons in reading frames 0, 1 and 2 of the influenza genome over time in human, avian, and swine lineages.

**Fig S3.**
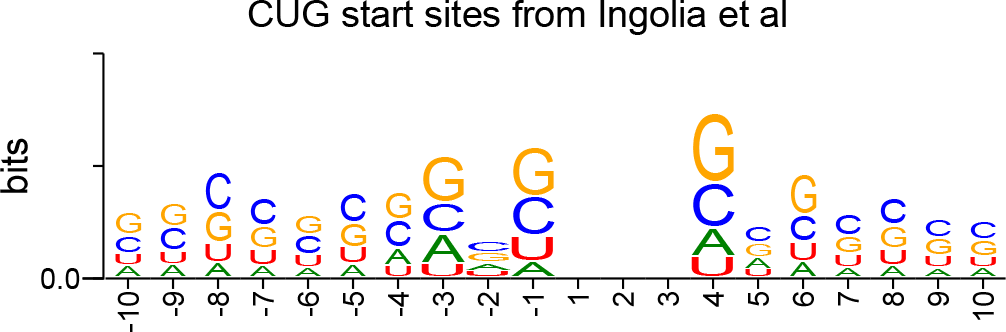
The consensus motif for CUG start codons. The consensus CUG start site motif was calculated using the Ingolia ribosome profiling dataset [30]. The most prominent feature of the consensus is a G at the +4 position.

**Fig S4.**
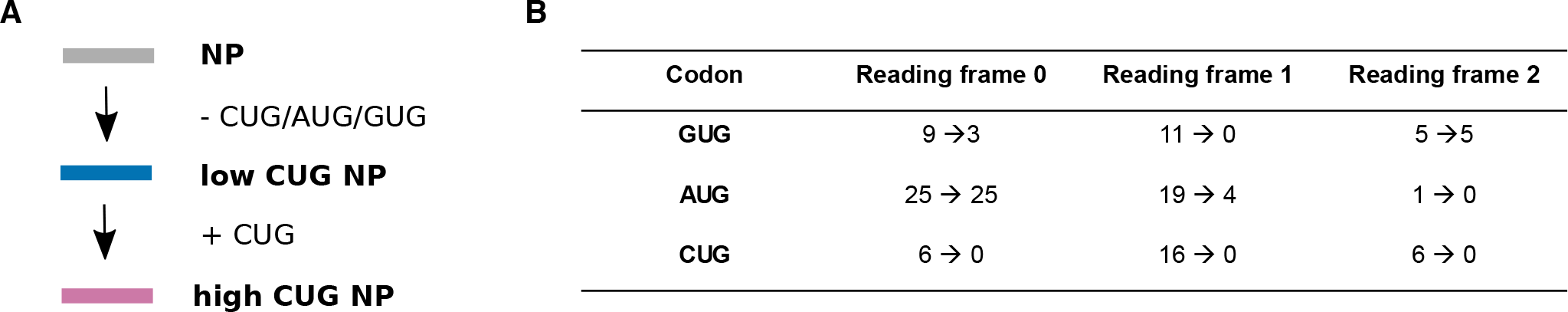
Summary of recoding influenza NP to generate low and high CUG NP. **(A)** Low CUG NP was generated by depleting wildtype PR8 NP of the common alternate start codons AUG, CUG, and GTG in all reading frames under constraints explained in Methods. High CUG NP was generated by adding 20 CUG codons into the low CUG NP background. All changes were synonymous with respect to reading frame 0. **(B)** Summary of the differences between wildtype PR8 NP (first number) and low CUG NP (second number) at the indicated codons.

**Fig S5.**
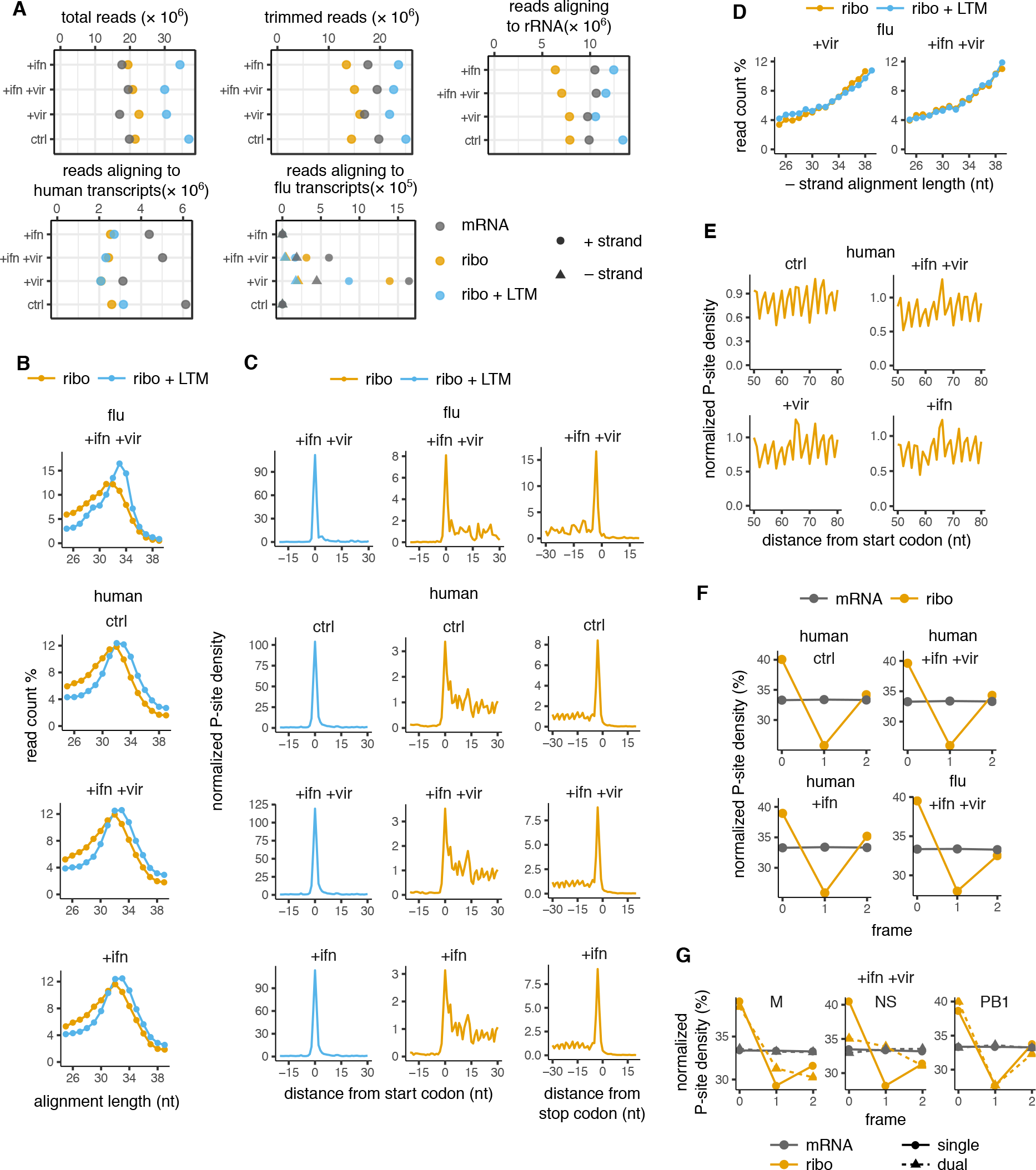
Summary of quality control steps for Ribo-seq data. **(A)** Number of input, trimmed, rRNA-aligned, and human/influenza transcript-aligned reads for assays and samples shown in Fig. 2. Number of reads aligning to + and − strands of influenza genome are shown separately. **(B)** Length distribution of viral (+ strand only) and human transcript aligned reads for Ribo-seq and Ribo-seq + LTM samples. **(C)** Metagene alignment of average P-site density around annotated start codons in viral and human transcripts for Ribo-seq + LTM and Ribo-seq samples. Right panels show the same for annotated stop codons for Ribo-seq samples. **(D)** Length distribution of viral (- strand only) genome aligned reads for Ribo-seq and Ribo-seq + LTM samples. **(E)** Metagene P-site density showing 3 nucleotide periodicity in a representative region of human transcripts for Ribo-seq samples. (F) Normalized P-site density in each of the reading frames of viral and human transcripts for RNA-seq and Ribo-seq samples. **(G)** Normalized P-site density in each of the reading frames of single- and dual-coded regions of M, NS, and PB1 segments of the influenza genome for RNA-seq and Ribo-seq samples for the +ifn +vir sample.

**Fig S6.**
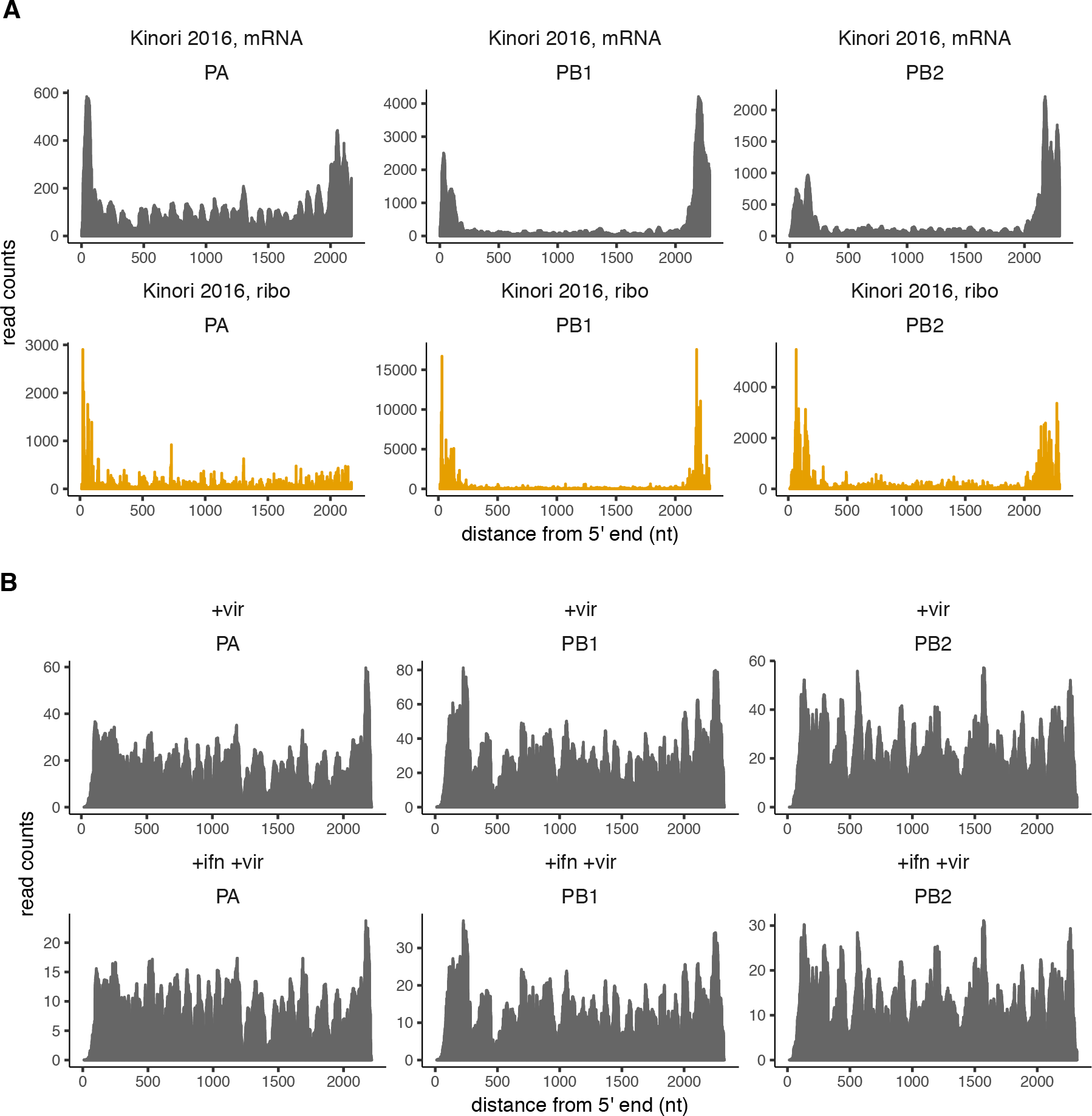
Read coverage of influenza polymerase segments reflects defective viral particles. **(A)** Distribution of RNA-seq and Ribo-seq read density along the polymerase segments of influenza. Raw data from [27]. **(B)** Distribution of RNA-seq read density along the polymerase segments of influenza in our data. See Fig. 3 for corresponding Ribo-seq density distribution. Defective viral particles are often characterized by the accumulation of large internal deletions in the polymerase segments. The large drop in coverage in panel A is consistent with the virus used in [27] containing a high burden of defective viral particles.

**Fig S7.**
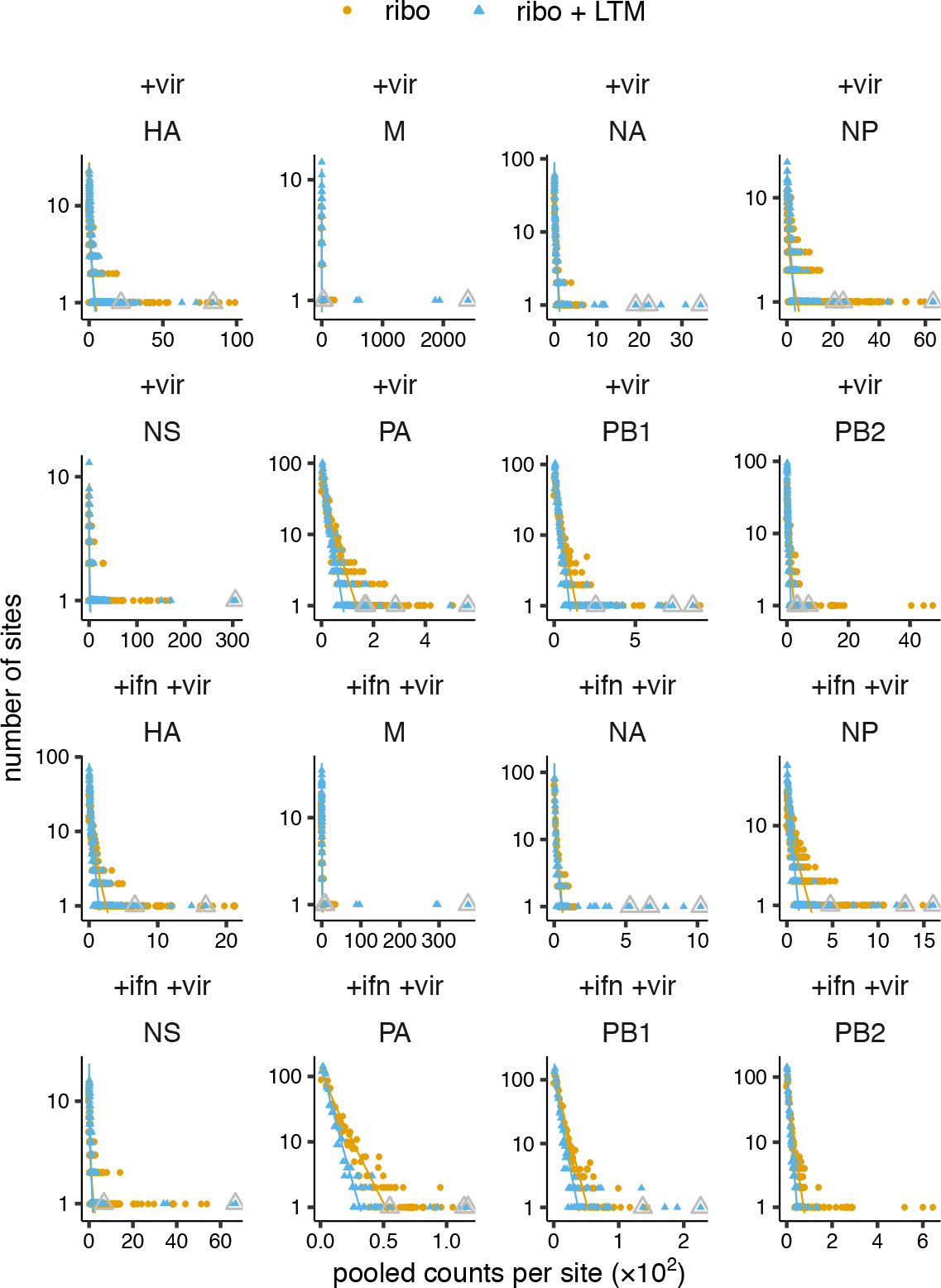
Background distribution fits for each influenza genome segment. The background Ribo-seq and Ribo-seq + LTM counts for each influenza genome segment were fit to separate zero-truncated negative binomial distributions (shown as lines). The final called TIS are indicated by grey triangles.

**Fig S8.**
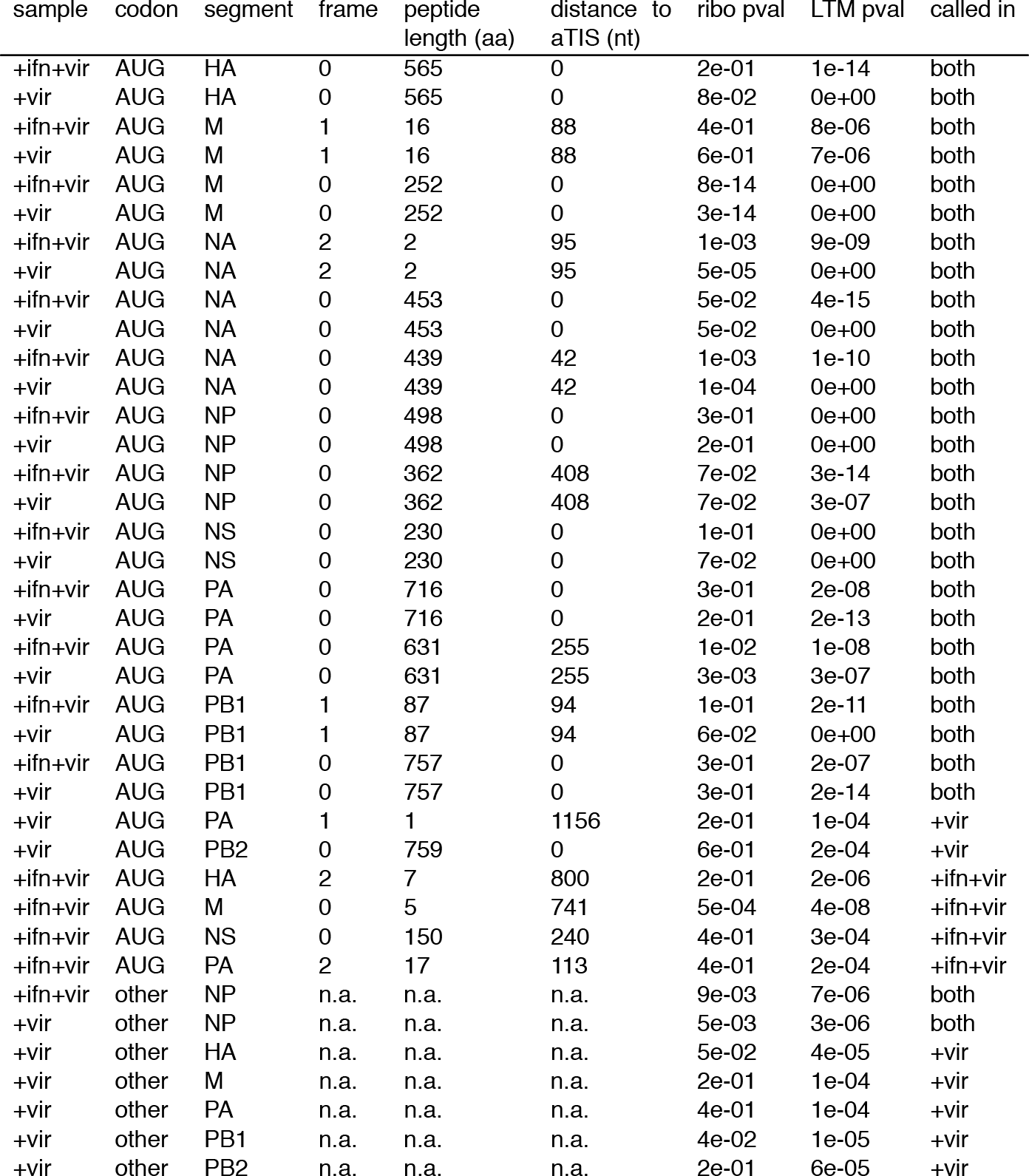
List of candidate start sites in the influenza genome.

**Fig S9.**
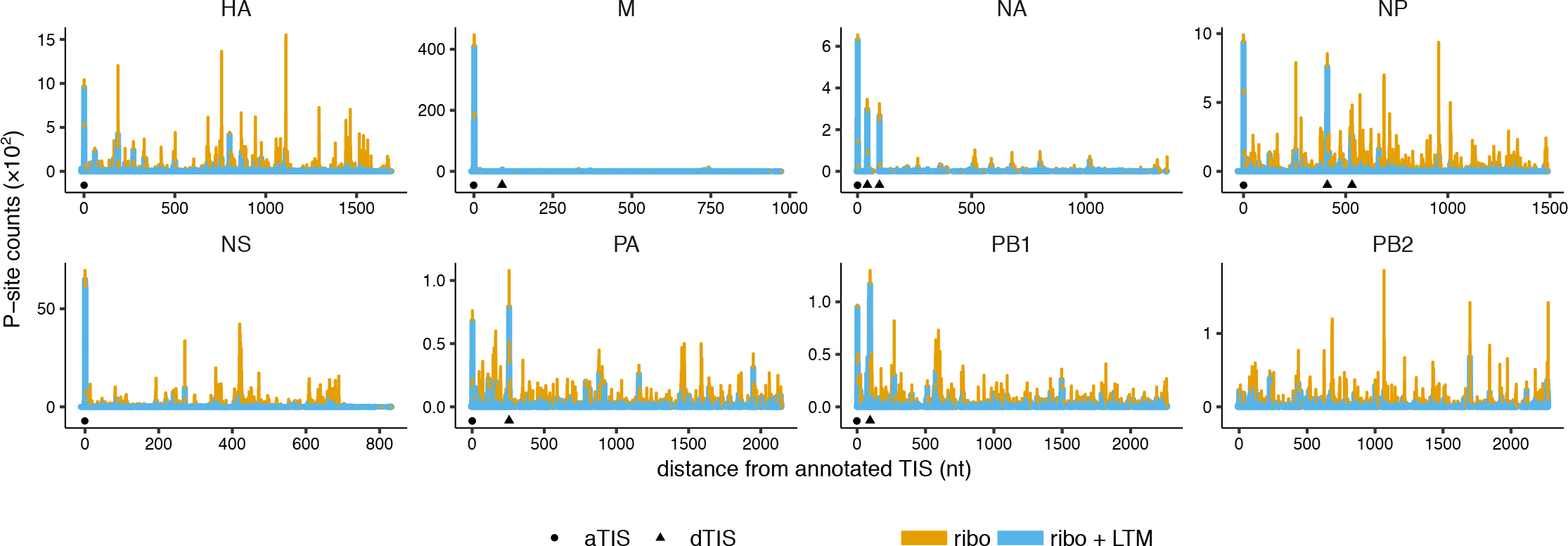
Ribo-seq coverage along influenza transcripts for +ifn +vir sample. P-site counts from Ribo-seq and Ribo-seq + LTM assays are shown for all 8 influenza genome segments for our +ifn +vir sample. The counts from the two assays are shown as stacked bar graphs for ease of comparison. The candidate annotated TIS (circle) and downstream TIS (triangle) shared between the +vir and +ifn +vir samples are indicated below the coverage plots.

**Fig S10.**
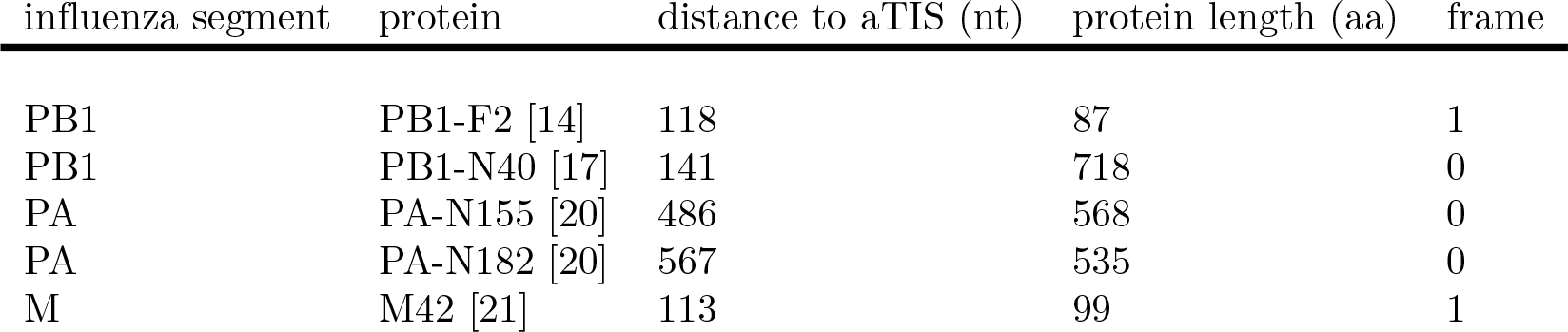
Summary of previously characterized alternate TIS in influenza. Summary of the literature concerning alternate TIS in influenza.

**Fig S11.**
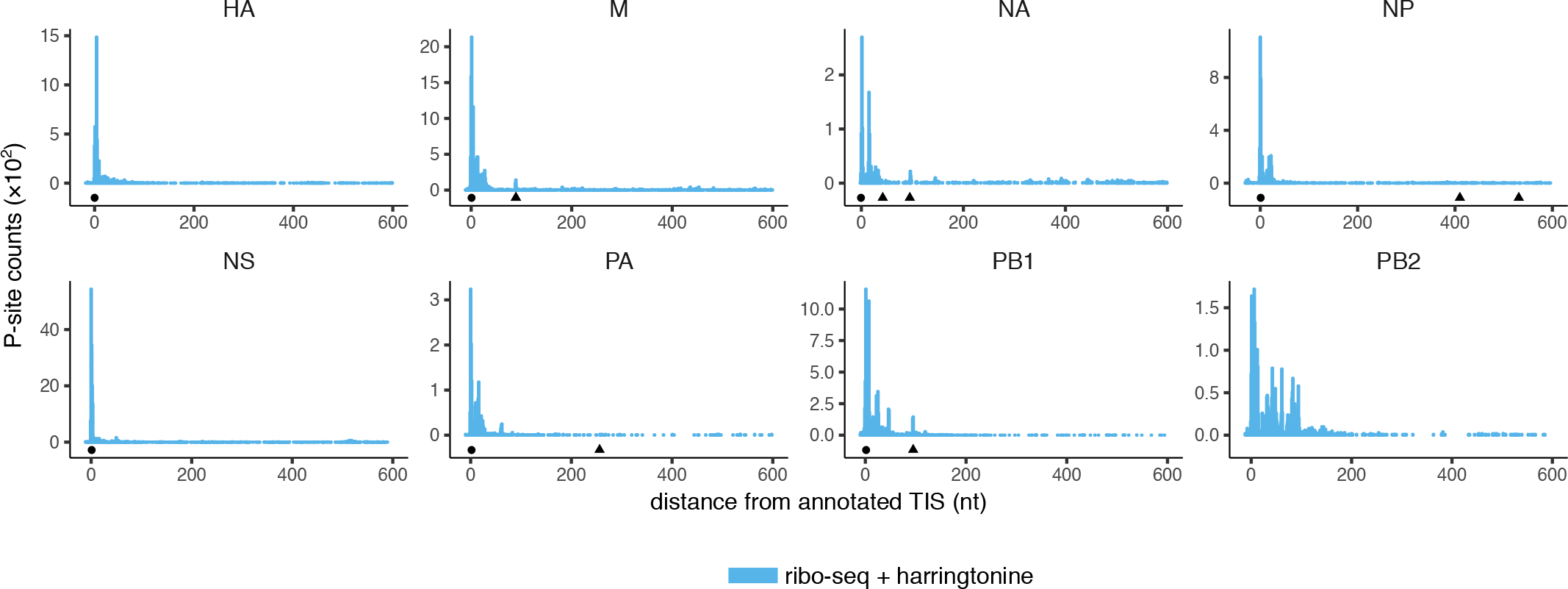
Peaks in Harringtonine-treated influenza samples from [27]. The P-site count coverage from [27] is overlaid with the 14 putative TIS identified in both of our +vir and +ifn +vir samples. Only the first 600 nucleotides of each gene sequence is shown.

**Fig S12.**
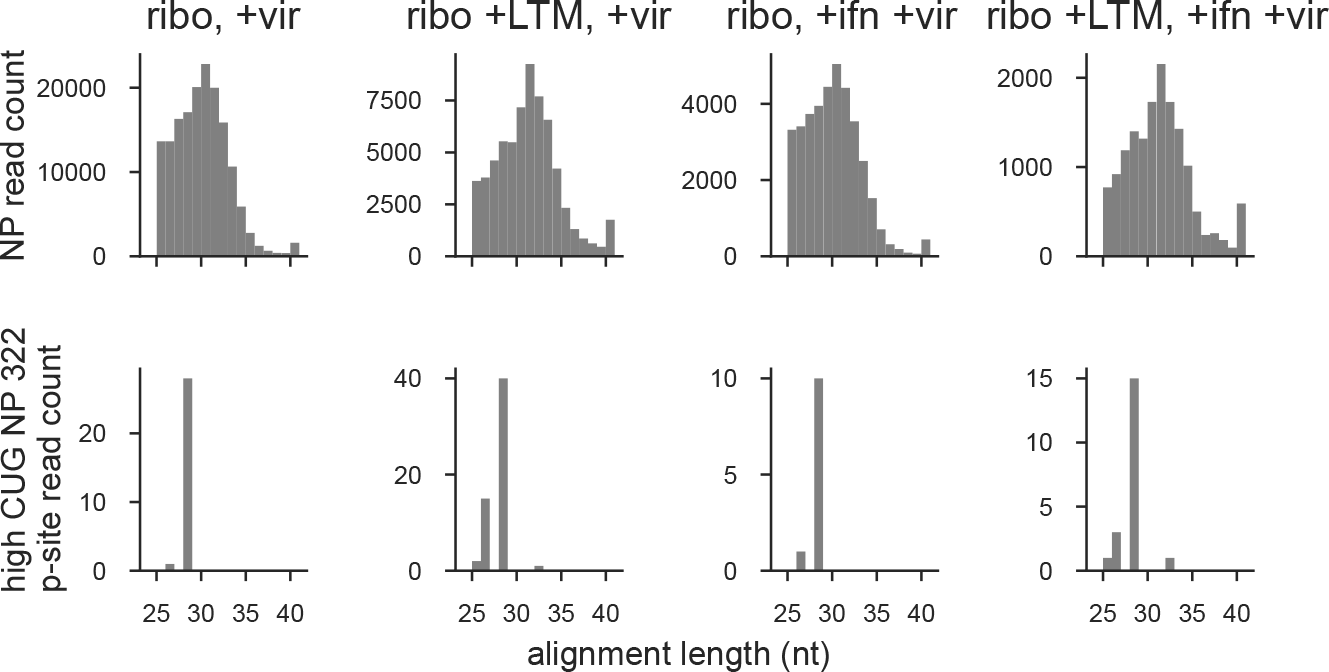
Ribo-seq alignment length for position 322 of high CUG NP. Distribution of alignment lengths for all NP reads and those with reads with P-site at position 322 of high CUG NP.

**Fig S13.**
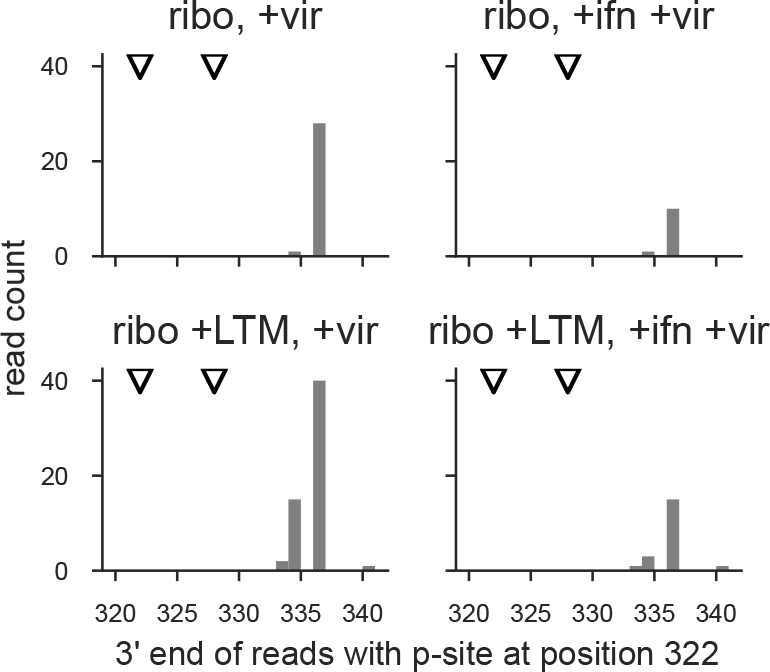
3′ end for reads with P-site at nucleotide 322 of high CUG NP. For high CUG NP reads with P-site position at nucleotide 322 the distribution of 3′ end of alignment is shown. Recoded CUG codons at nucleotides 322 and 328 are indicated with triangles.

**Fig S14.**
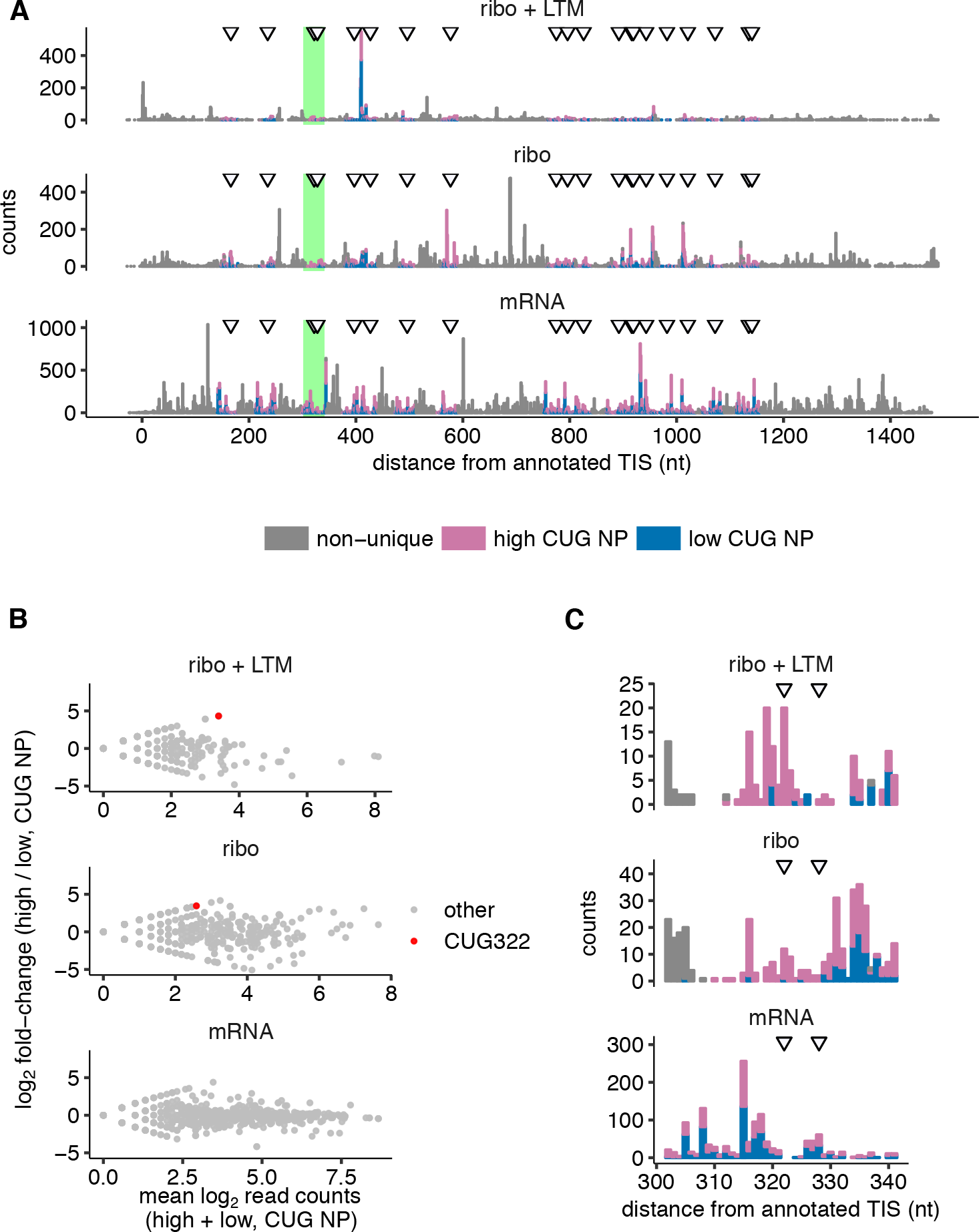
Translation initiation at CUG codons in influenza nucleoprotein in +ifn +vir sample. **(A)** Coverage of Ribo-seq + LTM, Ribo-seq, and RNA-seq reads that can be uniquely aligned to either the high CUG NP variant or the low CUG NP variant, and remaining non-unique reads. P-site counts are shown for Ribo-seq and Ribo-seq + LTM assays. 5′end counts are shown for RNA-Seq. Data are plotted as a stacked bar graph. Locations of the 20 CUG codons that are present in high CUG NP and synonymously mutated in low CUG NP are indicated by arrows. **(B)** The ratio of high CUG NP to low CUG NP coverage from A is plotted against their sum along the horizontal axis. There were no RNA-seq reads with counts at 322, so this point is not highlighted. **(C)** The green-highlighted region in A around the CUG322 codon is shown at greater horizontal magnification. See Fig. 4 for +vir sample.

**Fig S15.**
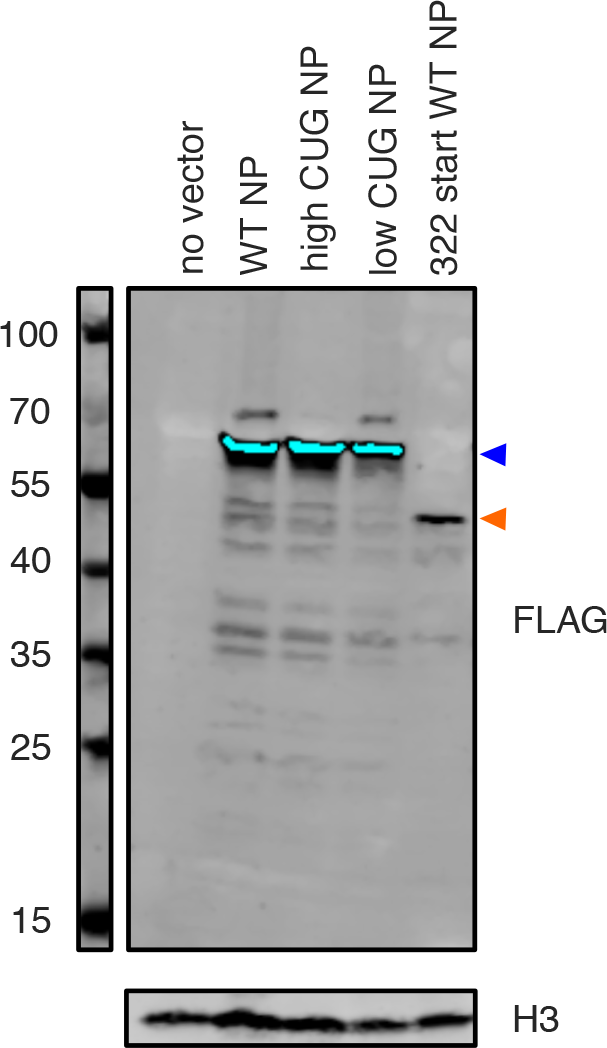
Western blot to examine possible initiation at CUG322 of high CUG NP. Western blot of 293T cells transfected with the indicated NP protein expression constructs. “322 start WT NP” is a size control construct that begins at nucleotide 322 of WT PR8 NP. Top panel: anti-Flag; bottom panel: anti-H3. Blue arrow corresponds to full length NP and orange arrow corresponds to expected size of NP fragment due to initiation at nucleotide 322. The blot was overexposed to sensitively detect truncated peptides, leading to saturation of the the full length NP band (shown in cyan).

**Fig S16.**
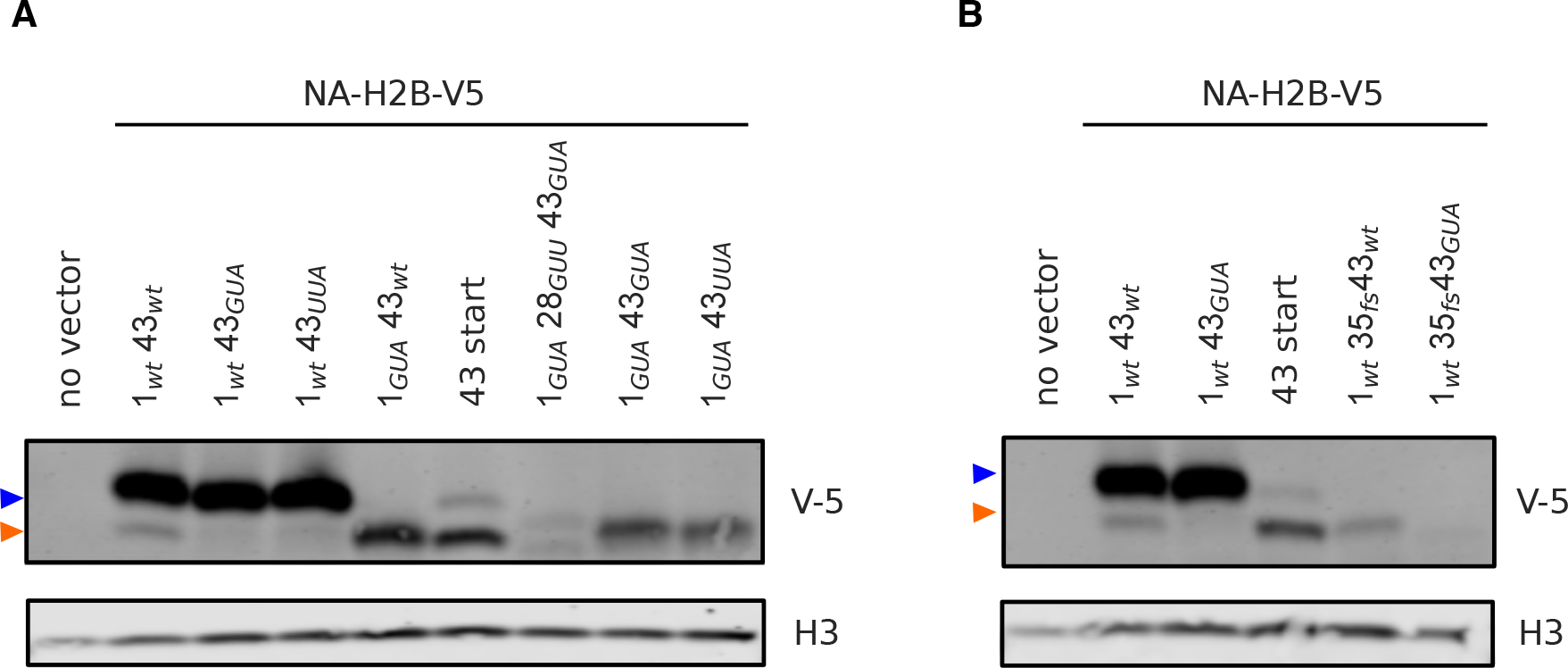
Western blot with additional constructs to examine initiation at NA43. **(A)** and **(B)** Western blot of 293T cells transfected with NA-H2B-V5 constructs with mutations at the canonical or downstream start site. “43 start” is a size control construct that begins at site 43 of NA. *1_GUA_28_GUU_43_GUA_* construct has a possible TIS (AUU codon) at position 28 mutated to GUU. *1_GUA_35_fs_43_GUA_* construct has a U inserted at coding nucleotide 35, such that any initiation 5′ to the insert should not be detectable with the V5 antibody. Top panel: anti-V5 (blue arrow corresponds to full length NA and orange arrow corresponds to NA43); bottom panel: anti-H3.

**Fig S17.**
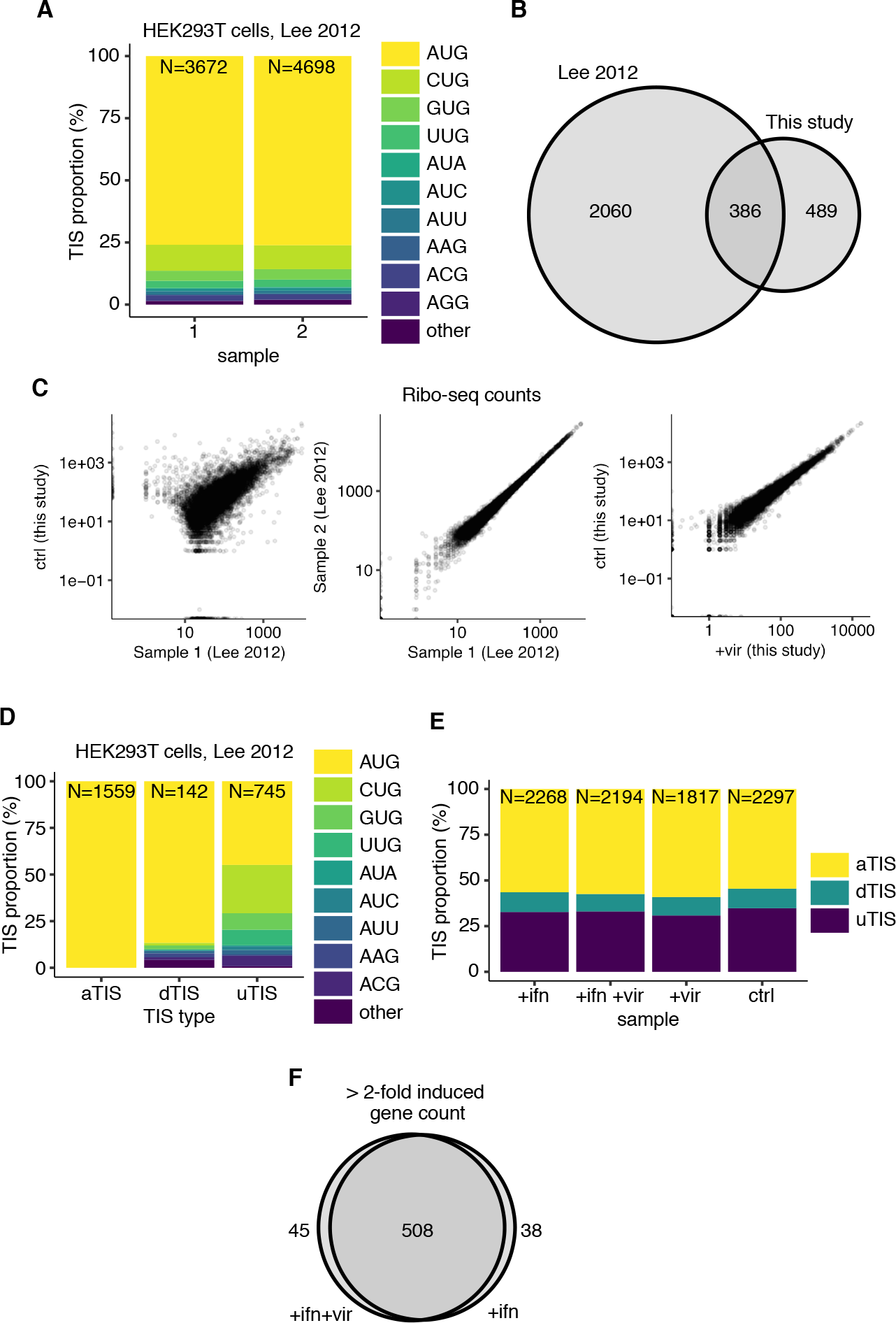
Statistics of called TIS in host transcripts. **(A)** Proportion of different near-cognate AUG codons (or other codons) overlapping with the called TIS in each of the two samples in [31]. *N* at the top of each bar indicates the total number of TIS called in each sample. **(B)** Overlap in high-confidence TIS between this study and Lee *et al.* [31]. High confidence TIS are the subset of TIS that are called across all samples in each study (2 in [31] and 4 in this study). **(C)** Comparison of Ribo-seq counts between samples in this study and from Lee 2012 [31]. Protein coding genes with at least 100 counts in one of the samples are plotted. **(D)** Proportion of different near-cognate AUG codons (or other codons) among the high-confidence TIS called in [31], stratified by TIS type. *N* at the top of each bar indicates the total number of high-confidence TIS of each type. **(E)** Proportion of different TIS types in each of the four samples used in this study. N at the top of each bar indicates the total number of TIS called in each sample. TIS not assigned to AUG or near-cognate AUG were excluded from this plot. **(F)** Overlap among the genes that are induced >2-fold upon either +ifn or +ifn +vir treatment with respect to the untreated sample. See Fig. 6 for definition of induced genes.

**S1 Table. Deep sequencing from NA43 competition.** Sequencing counts and ratios calculated for cell culture and mouse *1_wt_43_wt_* verses *1_wt_43_GUA_* and *1_wt_43_UUA_* virus competitions.

**S1 File. Influenza sequence alignments used for evolutionary analysis of CUG codons.** Alignments of protein-coding sequences of influenza PB2, PA, NP, M and NS to the A/Brevig Mission/1/1918 virus. Alignments were performed by appending the seven protein coding sequences together for each viral strain. PB2 is from position 1 to 2280, PA is from position 2281 to 4431, NP from position 4432 to 5928, M1 from position 5929 to 6687, M2 from position 6688 to 6981, NS1 from position 6982 to 7674, NS2 from position 7675 to 8040.

**S2 File. Influenza sequence alignments of NP used for generating low CUG PR8 NP and high CUG PR8 NP.** Alignments of protein-coding sequences of influenza NP.

**S3 File. Influenza sequence alignments of N1 NA.** Alignments of protein-coding sequences of influenza NA used for analysis of codon identity at position 43.

**S4 File. Influenza genome.** This file contains the influenza genome used for our ribosome profiling analysis, including low and high CUG PR8 NP sequences.

**S5 File. Influenza GTF.** This file contains annotations for influenza used for our ribosome profiling analysis.

